# Severe acute respiratory disease in American mink (*Neovison vison*) experimentally infected with SARS-CoV-2

**DOI:** 10.1101/2022.01.20.477164

**Authors:** Danielle R. Adney, Jamie Lovaglio, Jonathan E. Schulz, Claude Kwe Yinda, Victoria A. Avanzato, Elaine Haddock, Julia R. Port, Myndi G. Holbrook, Patrick W. Hanley, Greg Saturday, Dana Scott, Jessica R. Spengler, Cassandra Tansey, Caitlin M. Cossaboom, Natalie M. Wendling, Craig Martens, John Easley, Seng Wai Yap, Stephanie N. Seifert, Vincent J. Munster

## Abstract

An animal model that fully recapitulates severe COVID-19 presentation in humans has been a top priority since the discovery of SARS-CoV-2 in 2019. Although multiple animal models are available for mild to moderate clinical disease, a non-transgenic model that develops severe acute respiratory disease has not been described. Mink experimentally infected with SARS-CoV-2 developed severe acute respiratory disease, as evident by clinical respiratory disease, radiological, and histological changes. Virus was detected in nasal, oral, rectal, and fur swabs. Deep sequencing of SARS-CoV-2 from oral swabs and lung tissue samples showed repeated enrichment for a mutation in the gene encoding for nonstructural protein 6 in open reading frame 1a/1ab. Together, these data indicate that American mink develop clinical features characteristic of severe COVID19 and as such, are uniquely suited to test viral countermeasures.

**One Sentence Summary:** SARS-CoV-2 infected mink develop severe respiratory disease that recapitulates some components of severe acute respiratory disease, including ARDS.

## INTRODUCTION

Development of animal models has been a critical need since the emergence of SARS-CoV-2 in order to test vaccines and viral countermeasures (*1*). Multiple SARS-CoV-2 models of traditional laboratory species, including mice, hamsters, ferrets, and nonhuman primates, have been developed (*2–6*). However, these models either result in mild to moderate disease or experience distinctive viral dissemination due to altered angiotensin-converting enzyme 2 (ACE2) expression, specifically in the brain (*2,3,6*). An animal model that duplicates the severe acute disease spectrum of COVID-19 is still needed. Such a model would allow experimental research into the pathogenesis of severe COVID-19 and would facilitate the evaluation of therapeutic countermeasures in the context of severe disease.

Experimental infections of ferrets resulted in viral replication and transmission to naïve individuals, but very minimal to mild clinical signs of disease (*4,5*). However, this does not appear to be the case for all members of the Mustelidae family (*4,5,7-9*). Reports of farmed mink (*Neovison vison*) infected with SARS-CoV-2 emerged in the Netherlands in April 2020 (*10*). To date, at least 12 countries have reported outbreaks in farmed mink, as well as two reports describing positive feral or escaped mink (*11,12*). While many naturally infected mink exhibited mild to moderate clinical disease, a subset of these animals experienced an acute interstitial pneumonia that manifested with severe respiratory distress (*10,13*). Genomic surveillance in samples originating from mink in Denmark identified a series of changes in SARS-CoV-2 spike protein, known as the Cluster 5 variant (*14*). This mink-associated variant resulted in reduced neutralization with human convalescent sera *in vitro* (*14*). A high susceptibility to infection coupled with the public health risk of intra-host viral evolution prompted massive culls of an estimated 17 million mink on Danish farms (*15*).

Here, we show that experimentally infected mink develop a severe acute respiratory infection. After infection, progressive respiratory disease can be observed clinically, radiographically, and by histopathology. High amounts of viral RNA and infectious virus can be detected from the respiratory tract. Deep sequencing of SARS-CoV-2 genomes from oral swabs and lung tissue samples collected 3 days post-inoculation demonstrate rapid enrichment for a nonsynonymous mutation in the gene encoding for the nonstructural protein 6 (*nsp6*) in ORF1a in lung tissue samples. These data indicate the potential for rapid viral evolution in mink at the human-animal interface in a short timeframe. Together, these data suggest that the mink animal model recapitulates severe disease observed in hospitalized and fatal human cases of COVID-19 and could be useful to test countermeasures against severe COVID-19.

## RESULTS

### Mink ACE2 supports efficient entry of SARS-CoV-2

To evaluate the utility of the mink model for COVID-19, we compared human and mink ACE2 functional receptor entry using structural analysis and a vesicular stomatitis virus pseudotype entry assay (*16*). The sequence of mink (*Neovison vison*) ACE2 was obtained by sequencing from lung tissue. The obtained mink ACE2 sequence is 99.8% identical to the previously published *Neovison vison* ACE2 sequence (GenBank QPL1221.1). Two substitutions are observed at residues 231 and 613, in which the threonine and tyrosine in the previous sequence are replaced by a lysine and a cysteine, respectively. These residues are not located in the SARS-CoV-2 receptor binding domain (RBD)-ACE2 interface. Both American mink ACE2 sequences are 99% identical to the published European mink ACE2 sequence (*Mustela lutreola biedermanni*, GenBank QNC68911.1), and ∼83% identical to human ACE2 (*Homo sapiens*, GenBank BAB40370.1) (Figure 1A).

**Figure 1.**
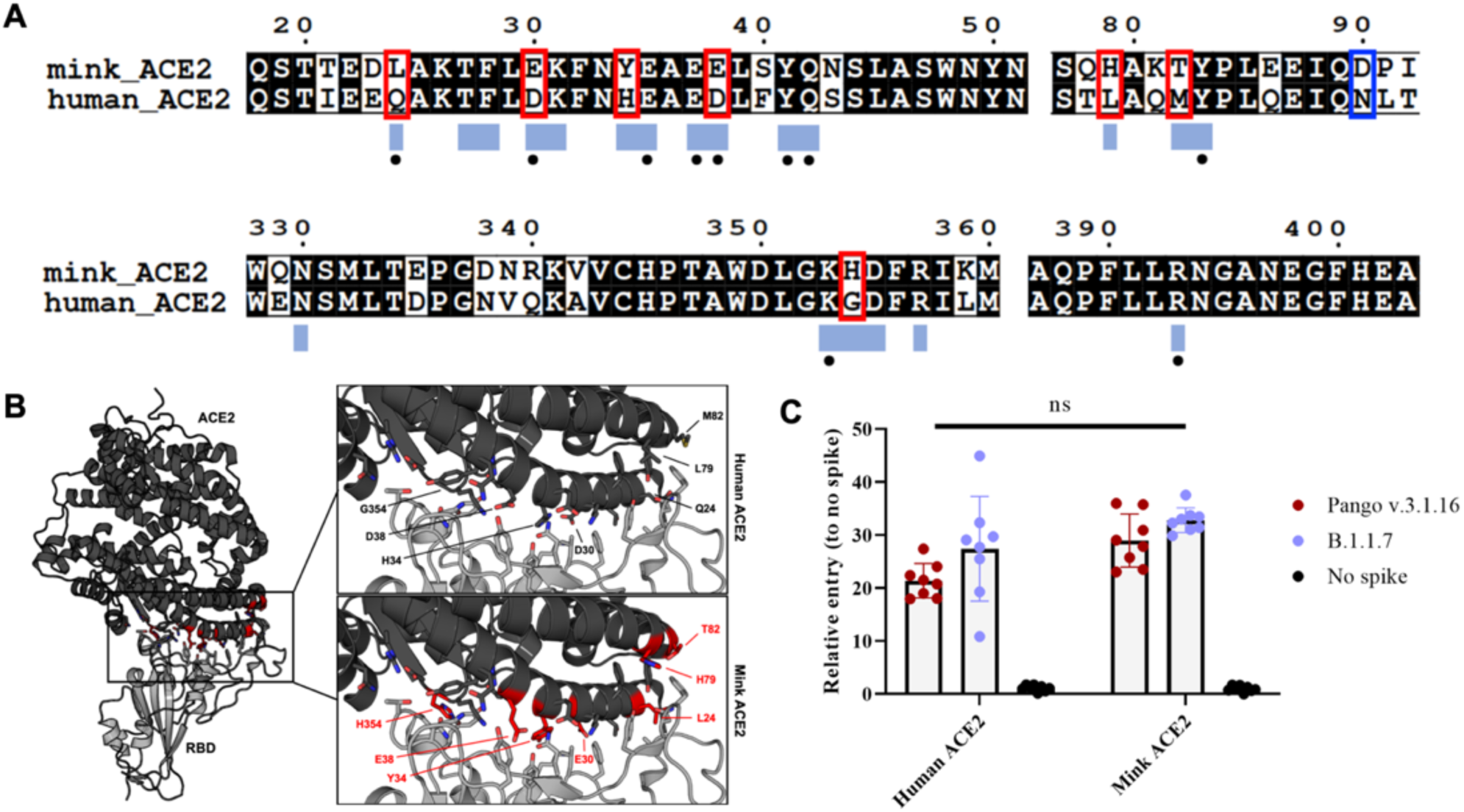
Functional SARS-CoV-2 entry analysis with human and mink ACE2. (A) An amino acid sequence alignment of ACE2 from the mink and human. Residues that participate in the SARS-CoV-2 RBD – ACE2 interaction are noted below the alignment by a blue box. Residues that participate in intermolecular hydrogen bonding or salt bridges are marked with a black dot. ACE2 residues that differ between mink and human within the interface are outlined with a red box. The substitution at residue 90 affecting an N-linked glycosylation site is noted with a blue box. (B) Differences between mink and human ACE2 are highlighted on the structure of the complex of SARS-CoV-2 RBD in gray bound to human ACE2 in black. Sidechains of the ACE2 and RBD residues that participate in the binding interaction are shown as sticks. The mutated residues are indicated by red. (C) SARS-CoV-2 spike pseudotype assay showing relative entry compared to no spike control in BHK cells expressing human or mink ACE2. Bars depict standard deviation.

To compare differences within the ACE2 interface with the SARS-CoV-2 spike receptor binding domain (RBD), the residues participating in the interaction, as described by Lan, *et al*. (*17*), were mapped onto an amino acid sequence alignment of ACE2 from American mink (*Neovison vison*), European mink (*Mustela lutreola biedermanni*, GenBank QNC68911.1), and humans (*Homo sapiens*, GenBank BAB40370.1)(*18*). The binding residues are 65% identical between mink and human ACE2, with seven of the 20 total interface residues differing in mink (Figure 1A). These residues are highlighted on the structure of SARS-CoV-2 RBD bound to human ACE2 to visualize these differences (Figure 1B). Consistent with a previous analysis, critical residues for interaction with the spike RBD, K31, Y41, and Y353 are conserved (*18*).

To investigate if the observed discrepancies between human and mink ACE2 translates to significant differences in spike entry, we directly compared the viral entry of VSV SARS-CoV-2 spike pseudotype particles on Baby Hamster Kidney fibroblasts (BHK cells) transfected with either human or mink ACE2. We observed significantly increased entry in mink ACE2 expressing cells compared to those expressing human ACE2 for the prototype WA1 lineage A SARS-CoV-2 spike (Figure 1C, alignment, entry data, p = 0.0307, 2-way ANOVA followed by Šídák’s multiple). However, the B.1.1.7 (Alpha) variant showed no difference (p = 0.1505, 2way ANOVA followed by Šídák’s multiple). Overall, the Alpha variant showed increased entry in both human and mink ACE2. Considering both variants together, there was no statistical difference in entry of the spikes to human and mink ACE2 (p = 0.5633, Two-tailed t-test) (Figure 1C).

We next determined the ACE2 expression in the respiratory tract of mink. ACE2 was multifocally detected in the respiratory olfactory epithelium and there were multifocal SARS-CoV-2 immunoreactive respiratory and olfactory epithelial cells (Supplemental Figure 1). ACE2 immunoreactivity was also detected in the lower respiratory bronchiolar epithelium and type I and type II pneumocytes (Supplemental Figure 1, D, E).

### Experimentally infected mink develop severe respiratory disease by 2 days post inoculation

Eleven adult farmed mink were inoculated intranasally and intratracheally with 10^5^ TCID_50_ of Alpha Variant, B.1.1.7 (hCoV-319 19/England/204820464/2020, EPI_ISL_683466). Due to the severity of clinical disease and respiratory distress, two animals reached end-point criteria and were euthanized the evening of 2 days post-inoculation (DPI). Eight animals reached end-point criteria on 3 DPI, and one animal recovered from severe disease and was euthanized on the predetermined experimental endpoint of 28 DPI.

Marked weight loss (up to 15%) was observed in all animals by 3 DPI (Figure 2A). In the animal that survived infection, bodyweight returned to baseline values by 14 DPI (Supplemental Figure 2A). Clinical signs were first detectable on 1 DPI in 5 of 11 (45 %) animals, with clinical signs observed in 9 of 11 (82 %) animals by 2 DPI and all remaining animals by 3 DPI. Signs of clinical disease included dull mentation, shivering, hunched or balled posture, lethargy, anorexia, increased respiratory effort, tachypnea, with occasional nasal discharge that included both epistaxis and serous discharge (Figure 2). Animals were examined on 1,3, 5, 7, 10, 14, 17, 21, and 28 DPI under anesthesia; 10 animals were clinically dehydrated by 3 DPI.

**Figure 2.**
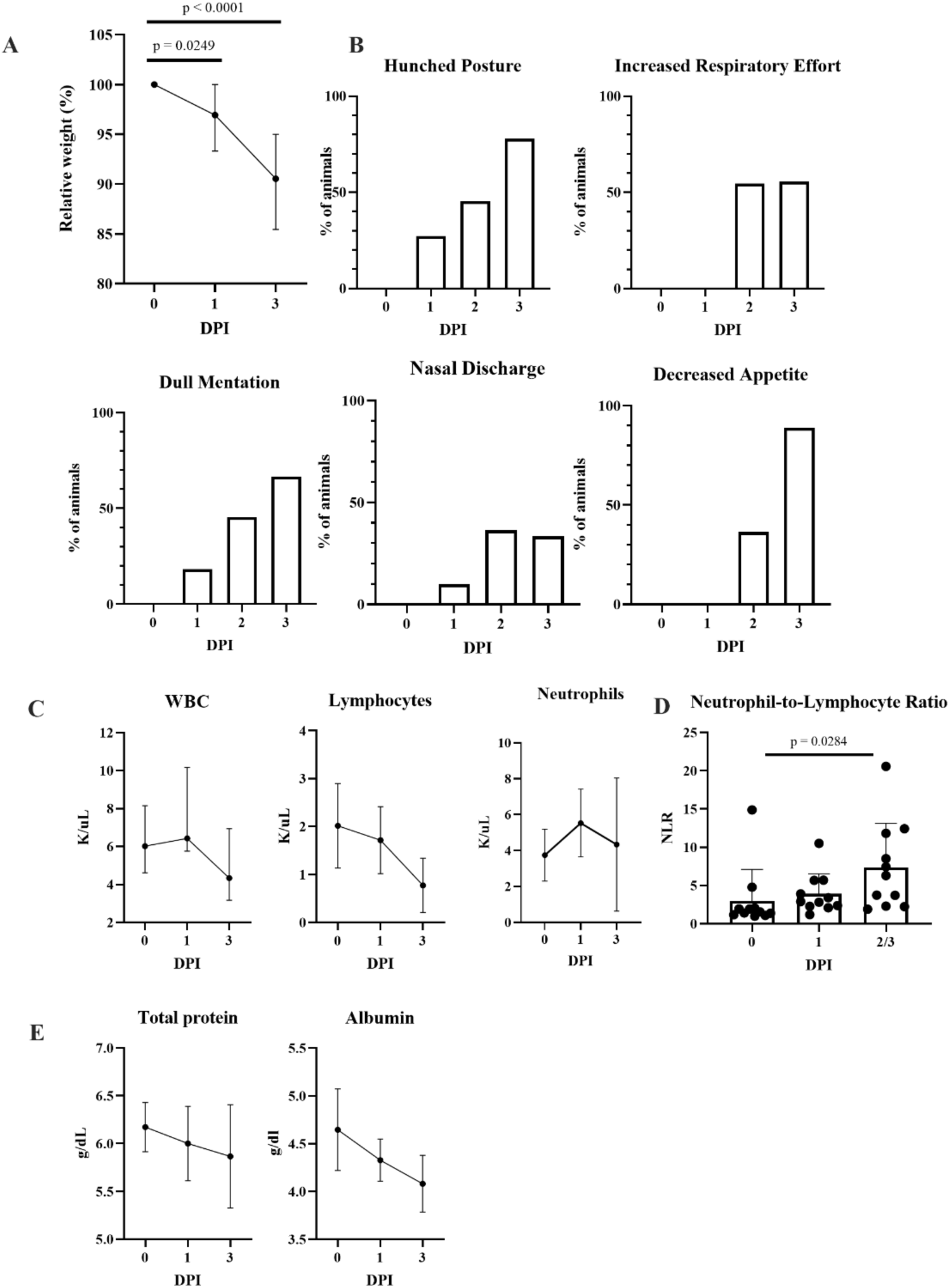
Features of acute respiratory disease in mink. (A) Percent of original body weight was collected during clinical exams on 1 and 3 DPI. Mink lost a significant amount of body weight on both 1 and 3 DPI (One-way ANOVA with Tukey’s multiple comparisons test). (B) Mink were assessed at least twice daily and evaluated for hunched posture, respiratory effort, mentation, nasal discharge, and appetite. (C) Complete blood count values collected after infection. The median with the 95% confidence interval (CI) are depicted. (D) Increased Neutrophil-to-Lymphocyte Ratio as determined from the complete blood count. Mean with standard deviation depicted, 2-way ANOVA with Tukey’s multiple comparisons test. (E) Selected blood chemistry values, median with 95% CI depicted.

Complete blood count (CBC) and complete chemistry panels were performed on blood samples collected at least one week prior to infection and at 0, 1, 3, 5, 7, 10, 14, 17, 21, and 28 DPI. At all time-points post infection, the CBC was unremarkable apart from a decreased white blood cell (WBC) count characterized by a mild lymphopenia that was most pronounced in the 9 remaining animals on 3 DPI (Figure 2C). The neutrophil-to-lymphocyte ratio was significantly increased at the terminal endpoint for clinically ill animals (Figure 2D). The single surviving animal had an elevated neutrophil-to-lymphocyte ratio (NLR) that peaked on 5 DPI as compared to baseline then quickly decreased (Supplemental Figure 2B). The blood chemistry panel was clinically unremarkable for all values except for a mild hypoproteinemia and hypoalbuminemia (Figure 2E).

### Progressive pulmonary infiltrates evident in pulmonary radiographs

Radiographic scores on 1 and 3 DPI were increased as compared to baseline values (Figure 3A) and indicated the presence of progressive pulmonary infiltrates consistent with viral pneumonia likely with concurrent non-cardiogenic pulmonary edema secondary to acute respiratory disease syndrome (ARDS)(Figure 3B). On 1 DPI, radiographic changes consistent with viral pneumonia were present in the thoracic radiographs of 5 (45%) of 11 mink. Of these 5, 4 had evidence of a mild-to-moderate ground glass/unstructured interstitial pattern and the remaining animal had a moderate-to-marked alveolar pattern affecting multiple lung lobes. Interestingly, this animal had progressive multifocal grade 3 and 4 pulmonary infiltrates at 2 DPI prior to euthanasia (Figure 3). At 3 DPI, 8 of the 9 remaining animals displayed disease progression that was characterized by increased severity and more extensive distribution of identified multifocal alveolar pattern (grade 3 to 4).

**Figure 3.**
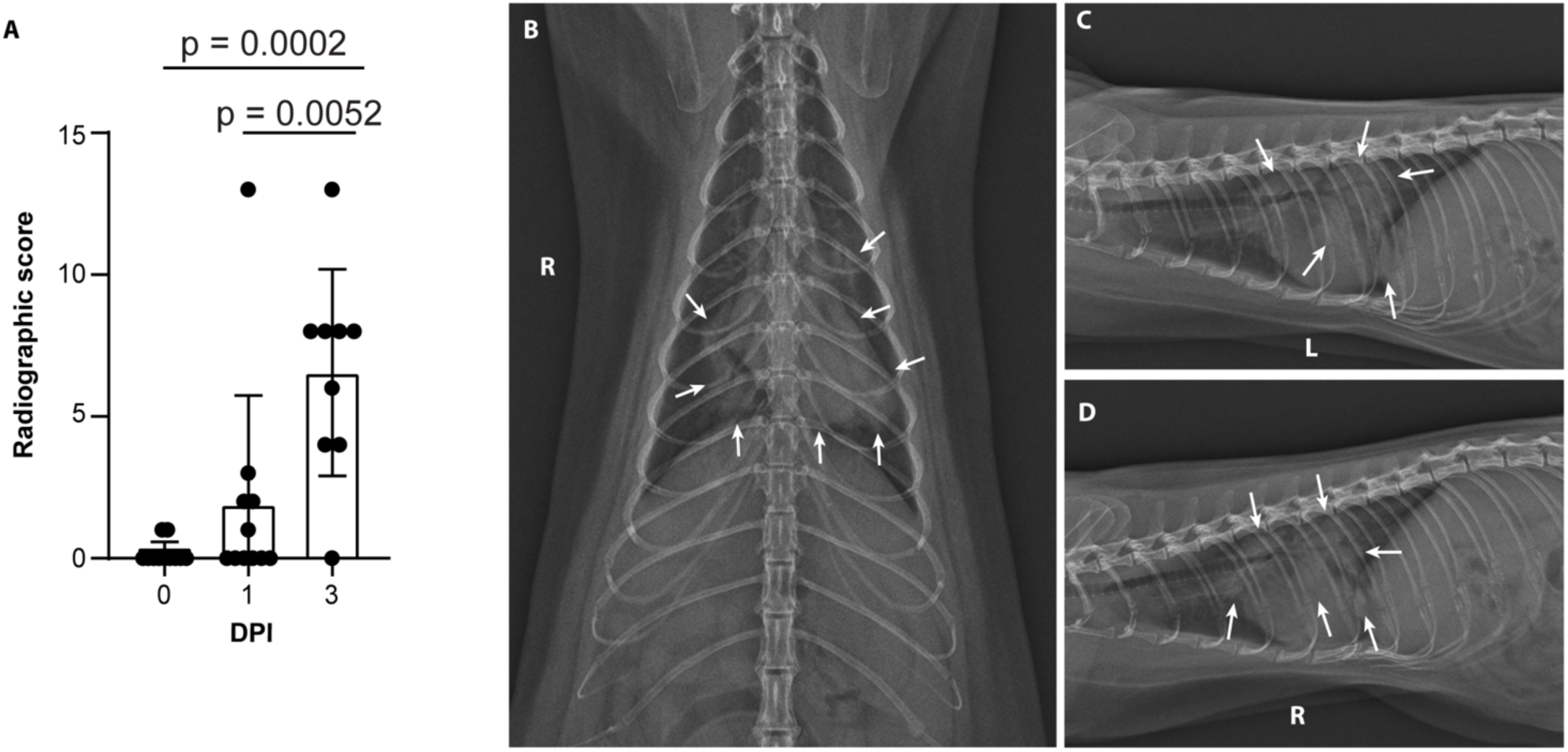
Severe radiological changes after infection with SARS-CoV-2. (A) Compiled radiographic scores. Bar graph depicts the mean with standard deviation and individuals, ordinary one-way ANOVA with Tukey’s multiple comparisons test. Radiographs demonstrate multifocal pulmonary infiltrates, most severe in the left and right caudal lung lobes depicted in the (B) dorsoventral radiograph (C) left lateral and (D) right lateral radiograph on evening of 2 DPI. Arrows depict grade 4 pulmonary disease in the left and right caudal lung lobes with grade 3 pulmonary disease in the right middle lung lobe and cranial subsegment of the left cranial lung lobe.

The remaining animal was monitored for resolution of disease over 28 days. Changes consistent with viral pneumonitis were first detected on 3 DPI, with the most severe changes noted on 5 DPI characterized by alveolar pattern in both caudal lung lobes. These changes began to resolve on 7 DPI with complete resolution noted on 14 DPI. (Supplemental Figure 3).

### Pathological changes in mink resemble severe human COVID-19 pulmonary damage and coagulopathy

Necropsy of all animals was performed immediately after euthanasia. Lung weight to body weight ratio was assessed to estimate the extent of pulmonary edema, and the ratio was significantly increased in infected animals compared to uninfected controls (Figure 4A). There were varying degrees of gross pulmonary pathology evident in all 10 animals euthanized on 2 or 3 DPI, with 100% of some lungs affected (Figure 4B). Grossly, lungs were hyperemic, and several animals had undergone pulmonary hepatization (Figure 4B).

**Figure 4.**
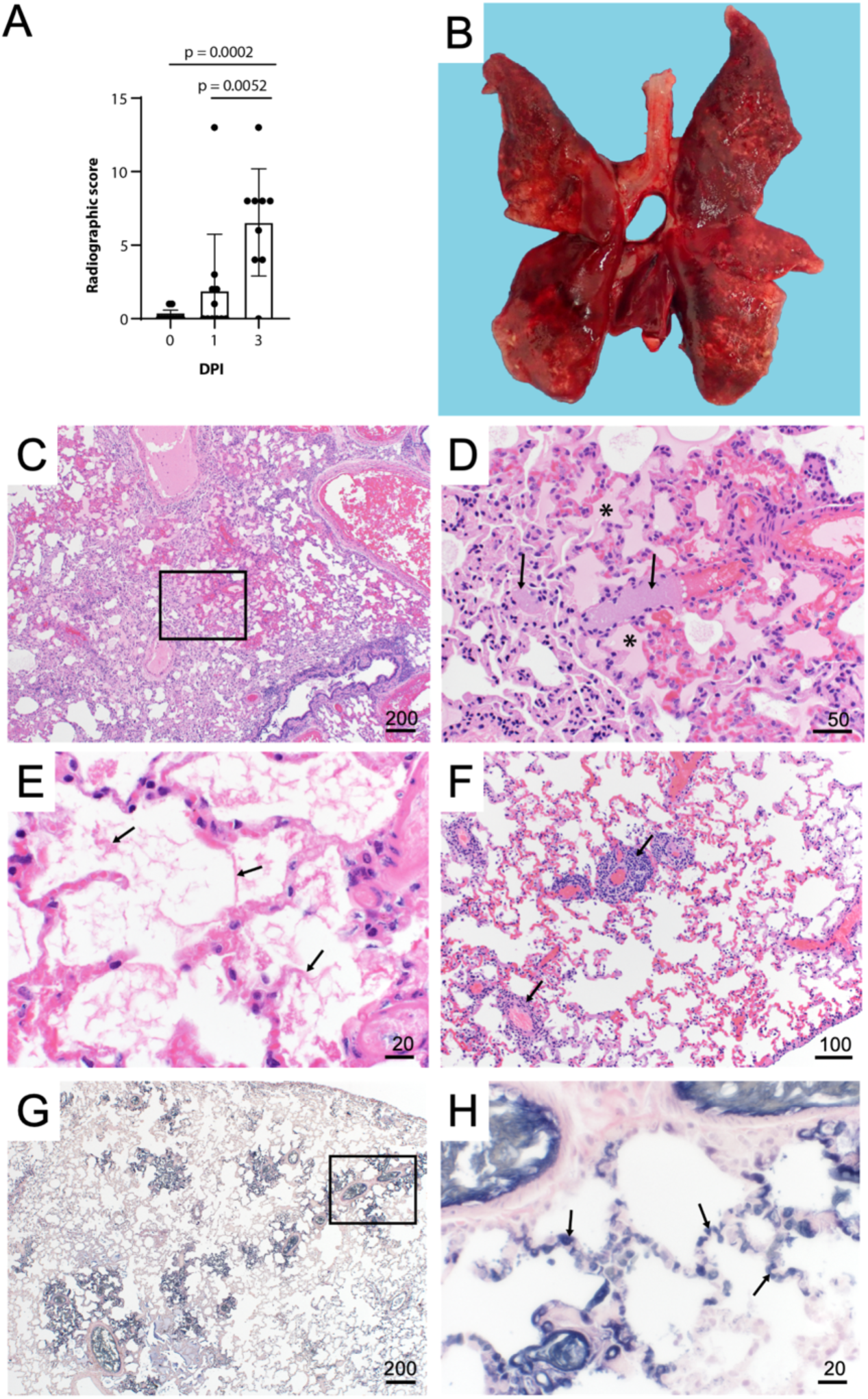
Pulmonary histopathology and immunohistochemistry. (A) Lung to body weight ratio, Mann-Whitney test. Graph depicts median with individuals. (B) Diffusely consolidated dark-mottled red lungs (C) Multifocal pulmonary congestion (Box) (D) Enlarged section of image C, vascular thrombi (arrows) alveolar edema (asterisks) (E) Alveolar fibrin (arrows) (F) Lymphoplasmacytic perivascular cuffing (arrows) (G) Pulmonary PTAH staining (black) (H) Enlarged section of image G, microthrombi (arrows). Scale bar expressed in μm in lower right corner of each image.

Histopathologic lesions associated with SARS-CoV-2 were restricted to the nasal turbinates and lungs of animals euthanized on 2 or 3 DPI. Nasal turbinates were characterized by a marked neutrophilic rhinitis with multifocal respiratory epithelial degeneration, necrosis, and loss. Nasal cavities were filled with an exudate composed of abundant neutrophilic and necrotic debris. There was rare neurotrophilic infiltration of olfactory epithelium (Figure 5). Pulmonary pathology was more severe in 3 of 10 animals euthanized on 2 or 3 DPI. Lesions consisted of moderate to marked vascular congestion (Figure 5, C, D) with thickening of the alveolar septa by edema, fibrin, and cellular infiltrate. Multifocal fibrin thrombi were identified in the vasculature of regions of congestion (Figure 5, E, F). Alveolar lumina often contained abundant edema fluid, fibrin, and increased number of alveolar macrophages. There was a moderate lymphoplasmacytic perivascular cuffing. Bronchial and bronchiolar epithelium was generally unaffected. The remaining seven animals had mild congestion, moderate lymphoplasmacytic perivascular cuffing with a mild increase in alveolar macrophages. SARS-CoV-2 antigen was observed predominately in pulmonary macrophages, although it was unclear if this was the result of replication or phagocytosis of viral antigen (Supplemental Figure 4, A, B). Multifocal SARS-CoV-2 antigen positivity was identified in bronchial epithelium and type I and II pneumocytes and bronchiolar epithelium (Supplemental Figure 5, C, D).

**Figure 5.**
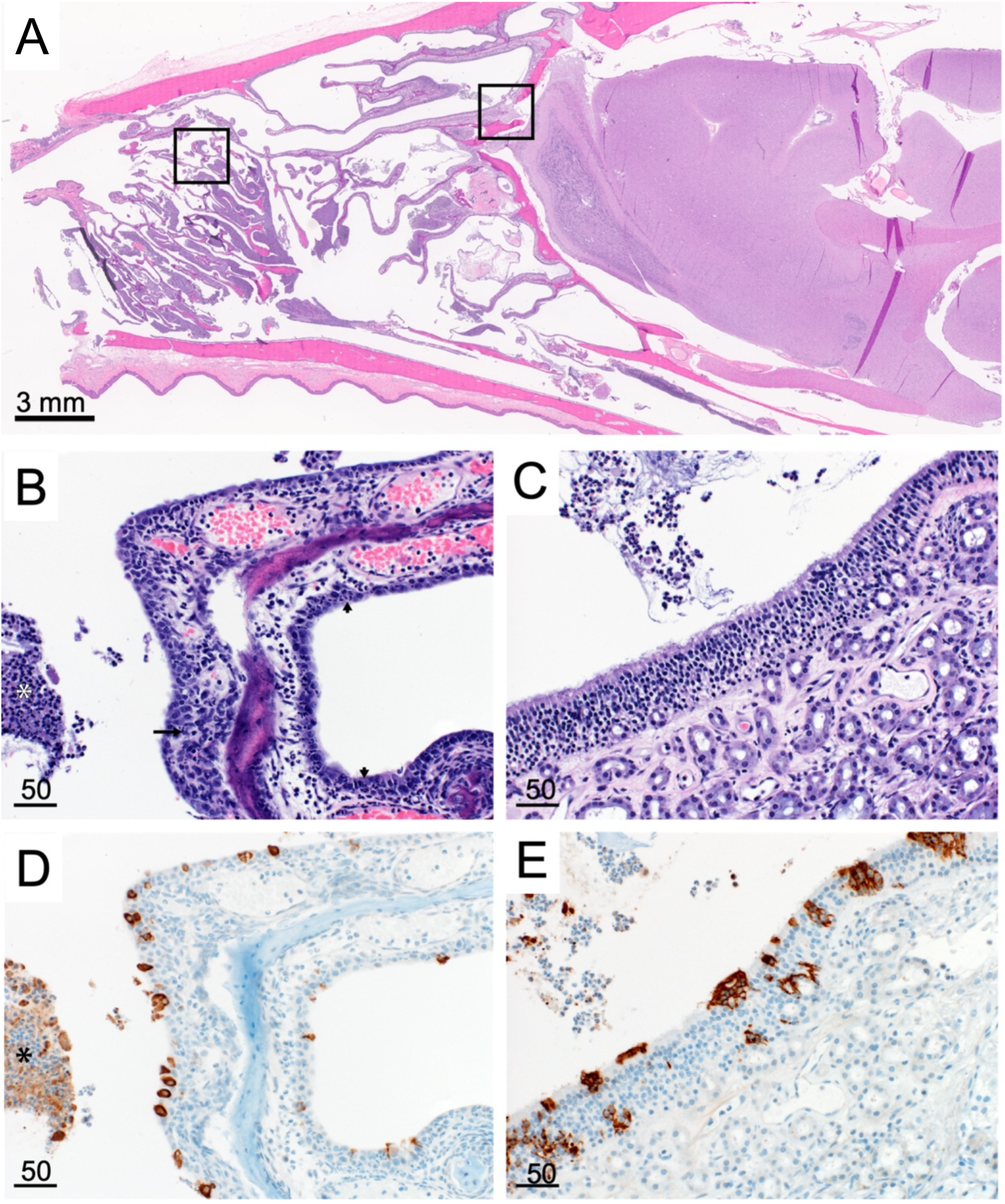
Nasal turbinate pathology and histopathology. (A) Sagittal section of skull, H&E (B) Respiratory epithelium neutrophilic infiltrates (small arrows) necrotic epithelium (long arrow) cellular exudate (asterisk) (C) Olfactory epithelium with cellular exudate (D) Respiratory epithelium SARS-CoV-2 IHC immunoreactivity (brown), immunoreactive cellular exudate (asterisk) (E) Olfactory epithelium SARS-CoV-2 IHC immunoreactivity and cellular exudate (brown). Scale bars expressed in μm unless indicated.

### Viral shedding is detected as early as 1 DPI in experimentally infected mink

Viral RNA was detected from all animals beginning on 1 DPI, with the highest viral RNA loads detected in oral and nasal swabs. In nasal, oral, and rectal swabs, both genomic (gRNA) RNA and sub-genomic RNA (sgRNA) were detected, with sgRNA as a marker for viral replication. Viral RNA was readily detected in both oral and nasal swabs (Figure 6, A, B). Viral RNA was also detected in rectal swabs collected from 4 of 11 (36 %) animals at 1 DPI and 3 of 9 (33%) animals at 3 DPI. (Figure 6C). Fur swabs were collected to estimate the risk of handlers and processers in the fur industry; genomic RNA was detected from most animals (Figure 4D). In the sole animal that survived until 28 DPI, viral RNA was detected in nasal swabs until 7 DPI and oral swabs until 10 DPI (Supplemental Figure 2C).

**Figure 6.**
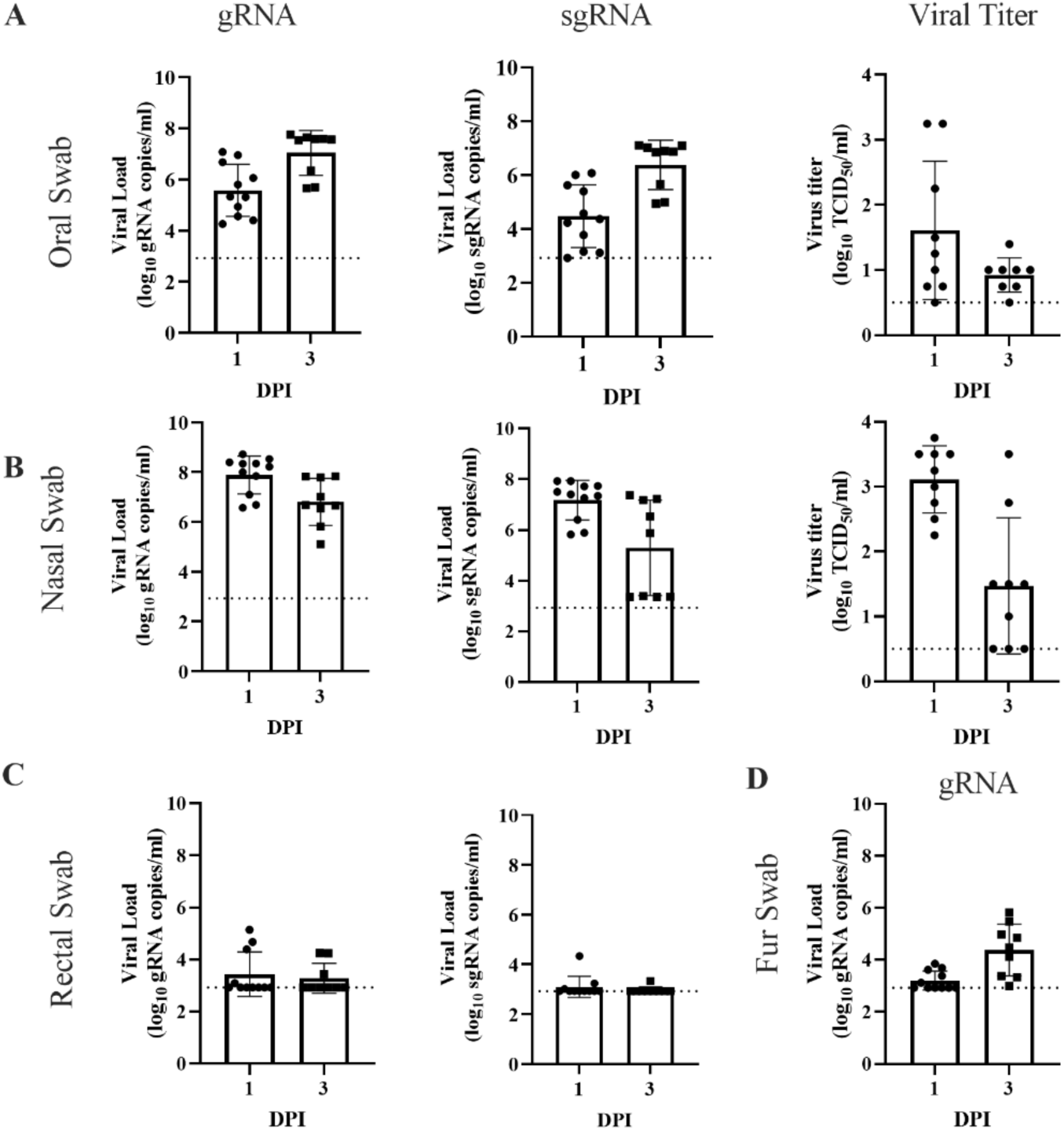
Viral shedding in infected mink. Oral (A), nasal (B), rectal (C), and fur (D) swabs were collected on 1 and 3 DPI and analyzed for genomic RNA, sub-genomic RNA, and infectious virus. Bar graphs depict the mean and standard deviation and individuals. Dotted line indicates the limit of detection.

Infectious virus was detected in most oral and nasal swabs at 1 (8 of 9, 89%) and 3 (7 of 8; 88%) DPI. Low amounts of infectious virus were detected in one rectal swab on 1 DPI (1.5 log10 TCID_50_/ml) and one fur swab on 3 DPI (0.75 log10 TCID_50_/ml). Infectious virus was detected in the surviving animal until 7 DPI in nasal swabs, and 10 DPI in oral swabs (Figure 6, A, B).

### High viral load in respiratory tract of SARS-CoV-2-infected mink

At necropsy, 37 tissues were collected from each animal and analyzed for the presence of both gRNA and sgRNA. gRNA and sgRNA was detected in the tissues from all animals necropsied on 2 or 3 DPI (Figure 7, Supplemental Figure 6). While viral RNA was detected in multiple organ systems, the highest viral loads were detected in respiratory tissues (Figure 7A, B). Within the respiratory tract, the highest viral loads were detected in the upper (nasal turbinate) and lower (all lung lobes) as compared to the mid-respiratory tract (trachea, right and left bronchus). In addition, high levels of viral RNA were detected in the frontal lobe, cerebellum, and brainstem (Supplemental Figure 6). Despite high levels of viral RNA, SARS-CoV-2 antigen was not observed in the olfactory bulb, cerebral cortex, or brainstem (Supplemental Figure 5). The respiratory tract was tested for the presence of infectious virus, with the majority of infectious virus found in the upper and lower respiratory tract (Figure 7C).

**Figure 7.**
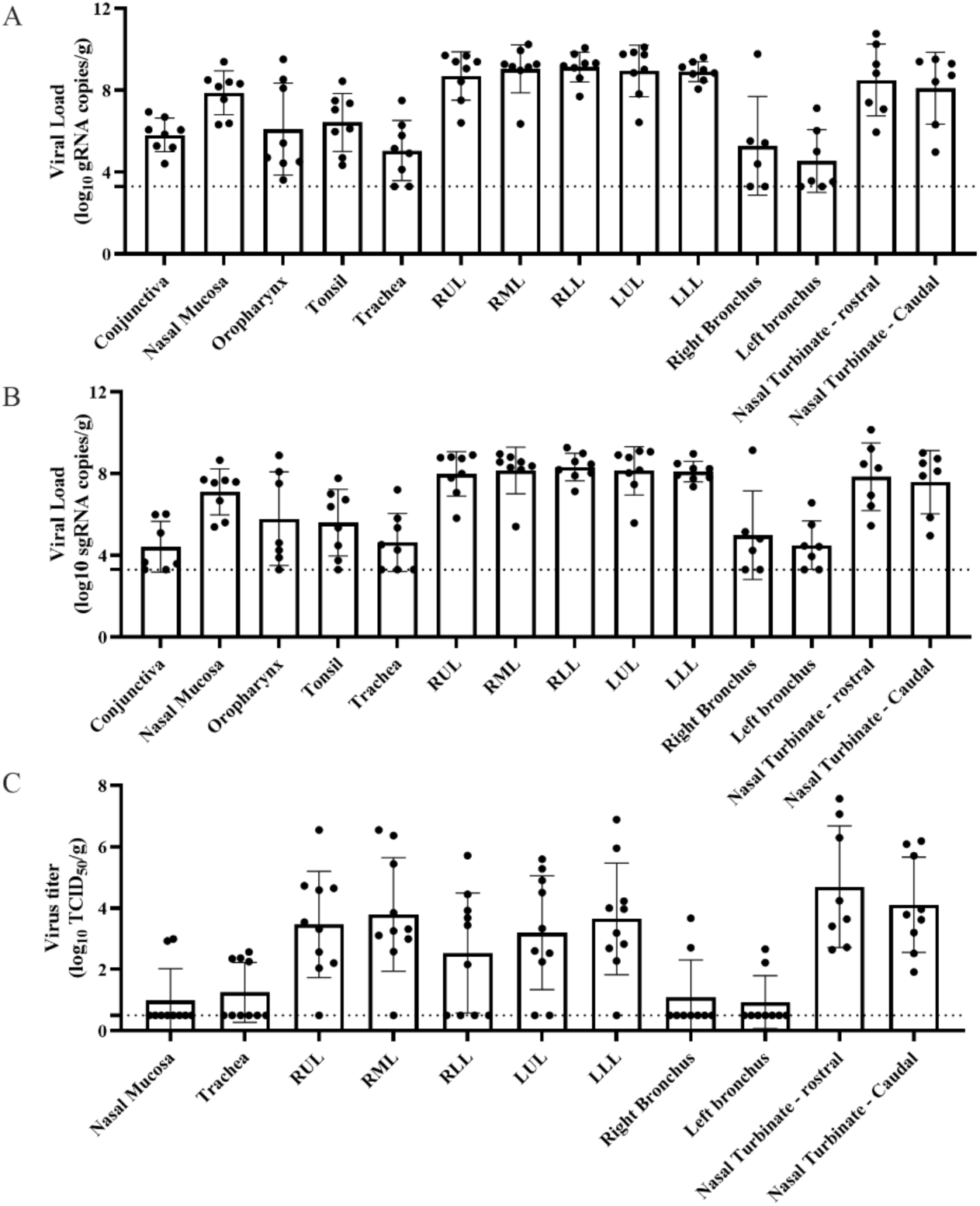
High levels of viral replication in the respiratory tract of infected mink. Tissues from animal euthanized on 3 DPI were analyzed for genomic (A), sub-genomic (B), and infectious virus (C). Bar graphs depict individuals, mean, and standard deviation. The dotted line depicts the limit of detection.

### Within-host evolution of SARS-CoV-2 in mink indicates potential for rapid adaptation

SARS-CoV-2 genomes were deep sequenced for 23 oral swabs and 10 lung tissue samples and the SARS-CoV-2 inoculum diluent used in the experimental challenge (Supplemental Table 1). The deep sequencing runs yielded an average of 88,686 reads mapped for each sample (Supplemental Table 1). One sample for which fewer than 50,000 reads were recovered was not used in subsequent analyses. Direct comparison of intrahost single nucleotide variants (iSNVs) detected at minor allele frequency thresholds of 3% and 5% showed a lack of concordance between technical replicates (Supplemental Figure 7), regardless of the SARS-CoV-2 genome copy number (Supplemental Figure 8A) or the number of sequencing reads mapped for the sample (Supplemental 8B). These results are consistent with findings in other deep viral genome sequencing efforts (*19,20*) and the need for tempered conclusions with the detection of low-level variants and small datasets. Therefore, we focus on changes in consensus sequence relative to the inoculum. Consensus sequences were largely unchanged from the SARS-CoV-2 inoculum sequence in all samples with the exception of a nonsynonymous mutation in the gene encoding nonstructural protein 6 (*nsp6*, L260F) which appeared enriched in the lung tissue samples of 5 mink relative to the inoculum and the oral swab samples (Figure 8). This mutation was identified eight times in 1064 available mink-associated SARS-CoV-2 genome sequences recovered from GISAID including a farmed mink in the USA (EPI_ISL_1014945) in Oct 2020, three mink the Netherlands (EPI_ISL_523102, 577749, and 523102) in May and Aug 2020, and four mink in Latvia (EPI_ISL_8514994, 8514995, 8514997, and 4548647, Supplemental Table 2) in July and Sept 2021.

**Figure 8.**
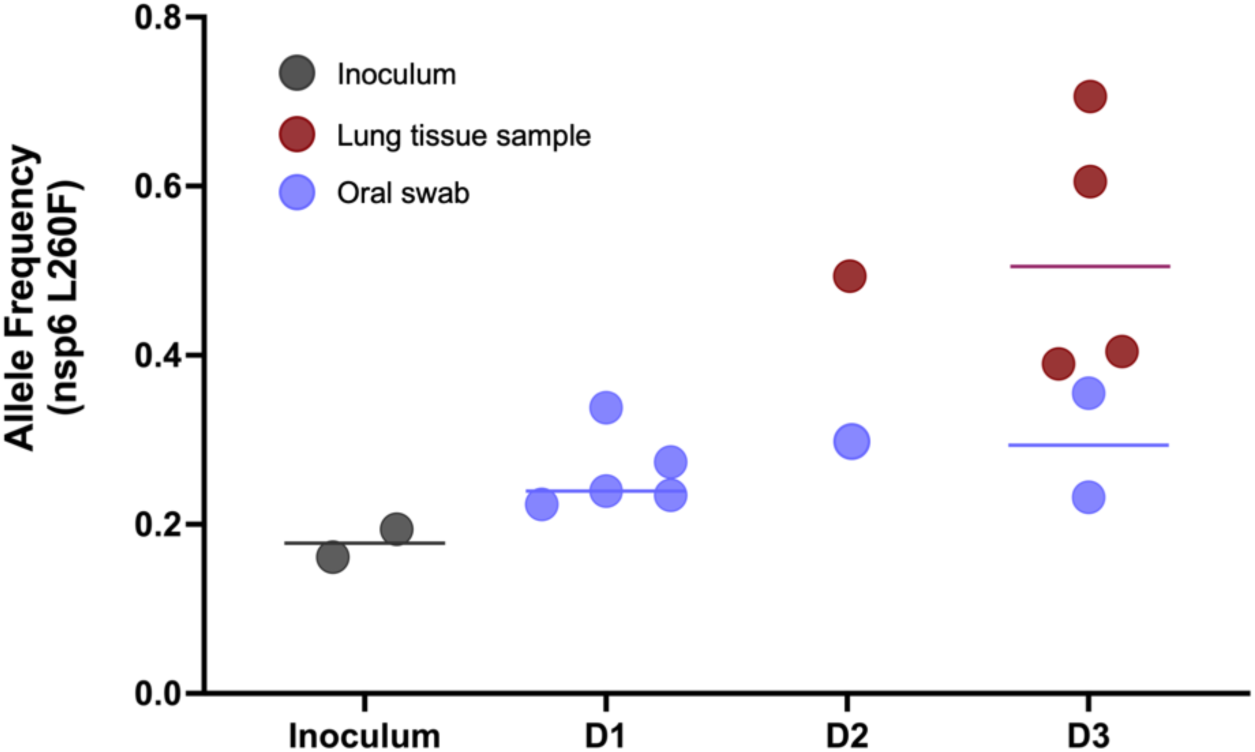
Enrichment of a nonsynonymous mutation (L260F) in the gene encoding for *nonstructural protein 6* in ORF1a of SARS-CoV-2 in oral swabs and lung tissue samples of five experimentally challenged mink. Deep sequencing of SARS-CoV-2 positive samples show rapid enrichment for L260F in *nsp6* in lung tissue samples but not the oral swabs for five of ten experimentally inoculated mink with SARS-CoV-2 genomic material detected in the lungs. Oral swab samples indicated by blue and lung tissue samples indicated by red, the line indicates group mean. Allele frequency for the L260F mutation is plotted on the y-axis.

### Seroconversion in surviving mink by 14 DPI

We analyzed serum for the development of a neutralizing antibody response. All animals had a titer of <20 prior to challenge and only the surviving animal developed a measurable neutralizing response. This animal seroconverted by 14 DPI with a peak neutralizing titer of 960, which decreased to a titer of 640 at euthanasia at 28 DPI.

## DISCUSSION

The continued emergence of SARS-CoV-2 variants of interest and variants of concern highlight the urgent need for animal models who consistently recapitulate the spectrum of disease in COVID-19 patients. Overall, the pseudotype entry data from this study demonstrates comparable spike entry between human and mink ACE2 confirming the suitability of mink for modeling SARS-CoV-2 infection and COVID-19 diseases.

Humans infected with SARS-CoV-2 present with a spectrum of clinical disease that ranges from asymptomatic infection to severe disease characterized by respiratory distress, sepsis, or multiorgan failure. Currently, an animal model for severe COVID-19 disease is not available (*1*). Most human infections are confined to the upper respiratory tract. Mild or early disease manifests with nonspecific symptoms that can include fatigue, fever, headache, loss of smell and taste, congestion, and fever (*21*). Progression into severe disease is typically presented as worsening respiratory disease, hypoxemia, and radiographic lesions, with end-point markers that can include coagulopathies, thromboembolism, acute kidney injury, and ARDS (*21–24*). Infected mink displayed clinical disease consistent with worsening human COVID-19 disease. Infected mink consistently demonstrated a greater degree of weight loss than that reported in nonhuman primates or hamsters in the days following infection. Increased respiratory effort and tachypnea in mink mark progression into severe COVID-19 disease. This study did not look at odor discrimination; however, neutrophilic infiltrate in olfactory epithelium could suggest a loss of smell that resulted in decreased appetite. Interestingly, while multiple field reports of fur farm outbreaks commonly report nasal discharge, this was not a consistent finding in these mink.

Features of complete blood counts of COVID-19 disease patients include leukopenia, lymphopenia, thrombocytopenia, and an increased NLR (*13,21,25,26*). While we were unable to rule out a stress leukogram resulting in increased NLR, this finding has been reported in ferrets infected with H5N1(*27*) and approximates critically ill human COVID-19 disease patients, where NLR can be used as a prognostic indicator (*25*). Interestingly, the NLR ratio for the single surviving animal was highest on 5 DPI, when the most severe changes were observed on thoracic radiographs. Mink displayed minimal to mild hypoproteinemia and hypoalbuminemia in the face of clinical dehydration, indicating a true hypoproteinemia. During acute disease, albumin can act as a negative phase protein and hypoalbuminemia has been associated with poor outcomes in COVID-19 patients (*28,29*).

One important hallmark of severe human COVID-19 disease is progression to ARDS. The Berlin Definition of ARDS addresses timing, thoracic imaging, the origin of thoracic edema, and the degree of hypoxemia (*23*). Our high-dose intratracheal inoculation likely contributed to the acute presentation of disease, and additional studies are necessary to better understand the course of SARS-CoV-2 infection in mink in relationship to dose, route of inoculation, and other emerging variants. Similar to the radiologic features described in humans, mink displayed bilateral ground glass opacities (*30*). However, unlike humans, these radiological features were not most severe in the gravitationally dependent regions (*30*). One theory explaining this atypical distribution is that the method of viral inoculation may have resulted in greater distribution in the caudal lung lobes as the virus was administered intratracheally in anesthetized subjects as opposed to a more passive inhalation of viral fomites. Additionally, a component of non-cardiogenic pulmonary edema, secondary to ARDS, may contribute to pulmonary infiltrates and is more commonly distributed in the caudal lung lobes. The hearts were radiographically and grossly normal, indicating the pulmonary changes were not likely due to cardiogenic pulmonary edema. Finally, this study was not able to evaluate the degree of hypoxemia, a critical step in diagnosing ARDS (*23*). Additional studies using advanced tools such as the flexiVent (SCIREQ, Emka Technologies Co., Sterling, VA, USA) and blood gas analysis are required to fully evaluate this model for clinical ARDS (*2,29,31,32*).

Histologically, diffuse alveolar disease (DAD) is an important finding in patients with severe COVID-19 disease and has not been regularly described in currently available animal models. Although not every mink had severe pulmonary pathology, all 10 animals euthanized on 2 or 3 DPI displayed pathology consistent with human COVID-19 disease (*24*). Animals with less severe histologic disease likely represent earlier stages in disease progression and follow up studies are required to better understand disease progression. The multifocal fibrin thrombi, cellular infiltrate, and resulting edema in the 3 mink with histologically severe disease likely reflects a coagulopathy, described in human patients with severe disease (*29,33*). Additional studies focusing on D-dimer, fibrinogen, and PT/aPTT are required to further tease out the pathogenesis in this model (*29*).

Outbreaks of COVID-19 on mink farms suggest the potential for novel SARS-CoV-2 variants to emerge in mink, with a high probability of spillback (*7-11,17*). This study provides experimental evidence of rapid enrichment in 5 of 10 mink for L260F in the gene encoding for nonstructural protein 6 in SARS-CoV-2 (Figure 8) which is hypothesized to affect viral autophagy and suppress the type I interferon response (*18,19*). This mutation has been identified in multiple COVID-19 outbreaks on mink farms in the Netherlands, Latvia, and the US. Interestingly, enrichment for the L260F mutation is most prominent in the lung tissue samples rather than the oral swabs, suggesting some tissue-specific tropism and likely a reduced probability of onward transmission. The repeated detection of the L260F mutation among COVID-19 outbreaks on mink farms through time and space supports that this mutation confers a selective advantage in mink and merits further study.

A pre-clinical model of severe COVID-19 disease is desperately needed to better evaluate SARS-CoV-2 vaccines and therapeutics. Current models demonstrate a reduction in viral titer and reduction of mild pathology, but no current model can recapitulate severe disease. In this study, we showed the utility of experimentally infected mink as a model for severe human COVID-19 disease. After infection, mink develop severe clinical disease associated with histological changes consistent with worsening human disease, making this new model the most translatable animal model available for severe COVID-19 disease.

## Materials and Methods

### Experimental Design

The objective of this study is to evaluate American mink as an animal model of severe COVID-19. SARS-CoV-2 infection in the American mink were determined through virological, histopathological, clinical, and radiographical analyses. Comparative analyses of functional SARS-CoV-2 entry using human and mink ACE2 were determined through a vesicular stomatitis virus pseudotyping assay. Deep viral genome sequencing was employed to study the intrahost evolutionary dynamics in the experimentally infected mink.

### Ethics statement

All animal experiments were approved by the Institutional Animal Care and Use Committee of Rocky Mountain Laboratories, NIH and carried out in an Association for Assessment and Accreditation of Laboratory Animal Care (AALAC) International accredited facility, according to the institution’s guidelines for animal use, following the guidelines and basic principles in the Guide for the Care and Use of Laboratory Animals, the Animal Welfare Act, United States Department of Agriculture and the United States Public Health Service Policy on Humane Care and Use of Laboratory Animals. The Institutional Biosafety Committee (IBC) approved work with infectious SARS-CoV-2 strains under BSL3 conditions. Sample inactivation was performed according to IBC-approved standard operating procedures for removal of specimens from high containment.

### Mink ACE2 Sequence and modeling

DNA was extracted from mink lung tissue using QIAamp DNA Tissue Kit according to the manufacturer. Mink ACE2 full length gene was amplified using long range PCR (LRPCR) amplification assay in two overlapping fragments using high-fidelity PrimeSTAR GXL DNA Polymerase (Takara Bio USA) as previously described (*34,35*). Briefly, 50 µL LRPCR master mix contained 0.2 µM of each primer (Supplementary Table 3), 1X PrimeSTAR GXL Buffer, 200 µM each deoxyribonucleotide triphosphate, 5 µL cDNA template, and 1.25 units of PrimeSTAR GXL DNA Polymerase (Takara Bio USA, Inc., San Jose, CA). The LRPCR mixture was incubated at 98°C for 2 minutes for the initial denaturation, followed by 4 cycles at 98°C for 10 seconds, 68°C for 15 seconds (−2°C per cycle), and 72°C for 10 minutes before an additional 26 cycles of 98°C for 10 seconds, 56°C for 15 seconds, and 72°C for 10 minutes. Sequencing libraries were generated using the TruSeq DNA PCR-Free library prep kit (Illumina Inc., San Diego, CA, USA) and sequenced on an Illumina MiSeq instrument at 2 x 151 paired-end reads. Reads were *de novo* assembled using SPAdes v. 3.13 (*36*). Sequence alignments between American mink (*Neovison vison*, sequence generated in this study) ACE2, European mink ACE2 (*Mustela lutreola biedermanni*, GenBank QNC68911.1), and human ACE2 (*Homo sapiens*, GenBank BAB40370.1) were generated using Multalin (*37*) and plotted using ESPript (*38*). Residues that participate in the SARS-CoV-2 RBD–ACE2 interaction, as described by Lan et al. (*17*), are noted below the alignment. The percent identity between the ACE2 sequences was calculated by Clustal Omega (*39*).

Structure analysis utilized the human ACE2 and SARS-CoV-2 RBD crystal structure, PDB ID 6M0J (*17*). Mutagenesis to show residues that differ in mink ACE2, and the alpha and delta variant RBD, was performed in COOT (*40*). The figures were generated using The Pymol Molecular Graphics System (https://www.schrodinger.com/pymol).

### Plasmids

The spike coding sequences for SARS-CoV-2 lineage B (hCoV-19/Denmark/DCGC-3024/2020, EPI_ISL_616802) were truncated by deleting 19 aa at the C-terminus. The S proteins with the 19 AA deletion of coronaviruses were previously reported to show increased efficiency regarding incorporation into virions of VSV (*41,42*). These sequences were codon optimized for human cells, then appended with a 5′ kozak expression sequence (GCCACC) and 3′ tetra-glycine linker followed by nucleotides encoding a FLAG-tag sequence (DYKDDDDK). These spike sequences were synthesized and cloned into pcDNA3.1^+^ (GenScript Biotech, Piscataway, NJ, USA). Mink ACE2 were synthesized and cloned into pcDNA3.1^+^ (GenScript Biotech, Piscataway, NJ, USA). All DNA constructs were verified by Sanger sequencing (ACGT).

### Pseudotype production and luciferase-based cell entry assay

Pseudotype production was carried out as described previously (*16*). Briefly, plates pre-coated with poly-L-lysine (Sigma-Aldrich, St. Louis, MO, USA) were seeded with 293T cells and transfected the following day with 1,200 ng of empty plasmid and 400 ng of plasmid encoding coronavirus spike or no-spike plasmid control (green fluorescent protein (GFP)). BHK cells were seeded in black 96-well plates and transfected the next day with 100 ng plasmid DNA encoding human or mink ACE2, using polyethylenimine (Polysciences, Inc., Warrington, PA, USA). After 24 hours, transfected cells were infected with VSVΔG seed particles pseudotyped with VSV-G, as previously described (*16,43*). After one hour of incubating with intermittent shaking at 37 °C, cells were washed four times and incubated in 2mL DMEM supplemented with 2% fetal bovine serum (FBS), penicillin/streptomycin and L-glutamine for 48 hours.

Supernatants were collected, centrifuged at 500xg for 5 minutes, aliquoted, and stored at −80 °C. BHK cells previously transfected with ACE2 plasmid of interest were inoculated with equivalent volumes of pseudotype stocks. Plates were then centrifuged at 1200xg at 4 °C for one hour and incubated overnight at 37 °C. Approximately 18-20 hours post-infection, Bright-Glo luciferase reagent (Promega Corp., Madison, WI, USA) was added to each well, 1:1, and luciferase was measured. Relative entry was calculated normalizing the relative light unit for each pseudotyped spike to the relative light unit average for the no-spike control.

### Animals

Seventeen apparently healthy adult farmed mink (*Neovison vison*) were used in this study: 11 were used for experimental infection and 6 were used as controls. All mink were pre-screened and negative for SARS-CoV-2 using a qRT-PCR, a pan-coronavirus assay (*44*), viral neutralization assay, and Aleutian disease using a lateral flow immunoassay (Scintilla Development Company LLC, Bath, Pennsylvania).

The animals used in the infection study consisted of 9 females and two males; intake female body weight range 1.04 kg – 1.47 kg, mean = 1.18 kg, male weights were 2.06 kg and 2.73. The females were approximately two years of age, the males were approximately one year of age. Upon arrival whole blood from all mink were screened for antibodies against SARS-CoV-2. Animals were single-housed in a climate-controlled room with a fixed light-dark cycle (12-hour light and 12-hour dark) for the duration of the experiment with access to food and water *ad libitum* with enrichment that included human interaction, commercial toys, music, and treats. All manipulations were done on anesthetized animals using Telazol (10-20 mg/kg administered subcutaneously).

### Animal study

Eleven animals were inoculated intratracheally (1.7 mL) and intranasally (0.15 mL per naris delivered using a MAD Nasal™ Mucosal Atomization Device (Teleflex, US) for a total dose of 10^5^ TCID_50_ delivered in 2 total mL. Animals were evaluated at least twice daily throughout the study. Clinical exams (including thoracic radiographs) were performed on 0, 1, 3, 5, 7, 10, 14, 17, 21, 28 DPI on anesthetized animals, during which the following parameters were assessed: bodyweight, body temperature, heart rate, respiratory rate, and radiographs. Clinical samples collected included nasal, oral, rectal, and fur swabs, and blood. Fur swabs were collected down the dorsal midline of the animal. Swabs were collected in 1mL of DMEM supplemented with 2% FBS, 1 mM L-glutamine, 50 U/ml penicillin, and 50 g/ml streptomycin.

### Radiographs

Ventrodorsal, left lateral, and right lateral thoracic radiographs were taken prior to clinical exams on 0, 1, 3, 5, 7, 10, 14, 17, 21, and 28 DPI with 0 DPI being performed prior to inoculation and serving as a baseline. Thoracic radiographs were taken immediately after animals were anesthetized and each lung lobe was evaluated by a board-certified veterinary radiologist as follows: 0 = normal lung, 1 = mild interstitial infiltrate, 2 = moderate to marked unstructured interstitial pattern, 3 = <25% alveolar pattern, 4 = >25% alveolar pattern.

### Clinical pathology

Hematology analysis was completed on a ProCyte Dx® (IDEXX Laboratories, Westbrook, ME, USA) and the following parameters were evaluated: red blood cells (RBC); hemoglobin (Hb); hematocrit (HCT); mean corpuscular volume (MCV); mean corpuscular hemoglobin (MCH); mean corpuscular hemoglobin concentration (MCHC); red cell distribution width (RDW); platelets; mean platelet volume (MPV); white blood cells (WBC); neutrophil count (absolute and percentage); lymphocyte count (absolute and percentage); monocyte count (absolute and percentage); eosinophil count (absolute and percentage); and basophil count (absolute and percentage). Serum chemistry analysis was completed on a VetScan VS2® Chemistry Analyzer (Abaxis, Union City CA) and the following parameters were evaluated: glucose; blood urea nitrogen (BUN); creatinine; calcium; albumin; total protein; alanine aminotransferase (ALT); aspartate aminotransferase (AST); alkaline phosphatase (ALP); total bilirubin; globulin; sodium; potassium; chloride and total carbon dioxide. Clinical pathology samples were evaluated by a board-certified clinical veterinarian.

### Histopathology

Histopathology and immunohistochemistry were performed on mink tissues. Tissues were fixed for a minimum of 7 days in 10% neutral-buffered formalin with 2 changes. Tissues were placed in cassettes and processed with a Sakura VIP-6 Tissue Tek, on a 12-hour automated schedule, using a graded series of ethanol, xylene, and PureAffin. Embedded tissues were sectioned at 5um and dried overnight at 42 degrees C prior to staining. The skulls were placed in Cancer Diagnostic acid free EDTA for 4 weeks and the solution was changed weekly.

Tissue sections were stained with hematoxylin and eosin (HE). The tissues were then processed for immunohistochemistry using the Discovery Ultra automated stainer (Roche Tissues Diagnostics) with a ChromoMap DAB kit (Roche Tissue Diagnostics cat #760-159). Specific anti-CoV immunoreactivity was detected using SARS-CoV-2 nucleocapsid antibody (GenScript Biotech, Piscataway, NJ, USA) at a 1:1000 dilution. The secondary antibody was the Vector Laboratories ImPress VR anti-rabbit IgG polymer (cat# MP-6401). To detect ACE-2, ACE-2 Antibody R&D Systems (catalog #AF933) was used at a 1:100 dilution with Vector Laboratories ImPress anti-goat IgG polymer (Cat #MP-7405) as a secondary antibody.

### Virus and cells

SARS-CoV-2 variant B.1.1.7 (hCoV-19/England/204820464/2020, EPI_ISL_683466; designated B.1.1.7 through the manuscript) was obtained from Public Health Agency England via BEI Resources. The obtained passage 2 material was propagated once in VeroE6 cells in DMEM supplemented with 2% FBS, 1mM L-glutamine, 50 U/ml penicillin, and 50 g/ml streptomycin. Mycoplasma testing was performed at regular intervals and no mycoplasma was detected. For sequencing from viral stocks, sequencing libraries were prepared using Stranded Total RNA Prep Ligation with Ribo-Zero Plus kit per manufacturer’s protocol (Illumina Inc., San Diego, CA, USA) and sequenced on an Illumina MiSeq at 2 x 150 base pair reads. Low level sequence variation in the stock of B.1.1.7 (nsp6/D165G/14%, nsp6/L257F/18% and nsp7/V11L/13%).

### RNA extraction and quantitative reverse-transcription polymerase chain reaction

RNA was extracted from nasal, oral, rectal, and fur swabs using the QiaAmp Viral RNA kit (Qiagen Sciences, Inc., Germantown, MD, USA) according to the manufacturer’s instructions and following high containment laboratory protocols. Tissue samples were homogenized and extracted using the RNeasy kit (Qiagen Sciences, Inc., Germantown, MD, USA) according to the manufacturer’s instructions and following high containment laboratory protocols. A viral sgRNA specific assay was used for the detection of viral RNA (*46*). Five μL of extracted RNA was tested with the Quantstudio 3 system (Thermofisher Scientific, Waltham, MA, USA) according to instructions from the manufacturer. A standard curve was generated during each run using SARS-CoV-2 standards containing a known number of genome copies.

### Viral titration

Tissue sections were weighed and homogenized in 1mL of DMEM. Virus titrations were performed by end point titration of 10-fold dilutions of swab media or tissue homogenates on VeroE6 cells in 96-well plates. When titrating tissue homogenate, the top 3 rows of cells were washed 2 times with DMEM prior to the addition of a final 100μL of DMEM. Cells were incubated at 37°C and 5% CO_2_. Cytopathic effect was read 6 days later.

### SARS-CoV-2 genome sequencing and within-host evolution

Oral swabs and lung tissue samples were collected into Trizol for RNA extraction. 200 μL of 1-Bromo-3-chloropropane (MilliporeSigma, St. Louis, MO, USA) was added to the Trizol/sample lysate, mixed, and centrifuged at 16,000 x *g* for 15 min at 4°C. RNA containing aqueous phase of 600 μl was collected from each sample and aqueous phase was combined with 600uL of RLT lysis buffer (Qiagen, Valencia, CA, USA) with 1% beta mercaptoethanol (MilliporeSigma, St. Louis, MO, USA). RNA was extracted using Qiagen AllPrep DNA/RNA 96-well system (Valencia, CA, USA). An additional on-column Dnase 1 treatment was performed during RNA extraction. All sample processing was performed using amplicon-free reagents and tools in aerosol resistant vials. RNA quality was analyzed using Agilent 2100 Bioanalyzer (Agilent Technologies, Santa Clara, CA, USA). RNA samples were quantitated by qRT-PCR targeting NSP5 using the AgPath-ID One-Step RT-PCR Buffer and Enzyme Mix (Life Technologies, Carlsbad, CA, USA). The reactions were carried out in 20 µL reactions using NSP5 forward primer (5’-CTGGCACAGACTTAGAAGGTAACTT-3’), reverse primer (5’TCGATTGAGAAACCACCTGTCT-3’), fluorescent probe (5’-6FAM-TTGACAGGCAAACAGCACAAGCAG-BHQ1-3’) (Biosearch Technologies, Novato, CA, USA). The QPCR reactions were carried out at 50 °C for 10 minutes, 95 °C for 10 minutes, 55 cycles of 95 °C for 15 seconds and 60 °C for 45 seconds. Data was analyzed using ABI 7900HT version 2.4 sequence detection system software (Thermofisher Scientific, Waltham, MA, USA) and SARS-CoV-2 genome copy (equivalent/mL) numbers were determined by absolute quantitation method. Next generation libraries were generated using the TruSeq DNA PCR Free Nano kit (Illumina, Inc., San Diego, CA, USA) and the ARTIC multiplex PCR genome amplification protocol with the V3 primer scheme (www.protocols.io/view/ncov-2019-sequencing-protocol-bbmuik6w) and libraries were sequenced on an Illumina MiSeq at 2 x 250 paired-end reads. The ARTIC multiplex PCR SARS-CoV-2 genome amplification protocol has been widely used in viral genome sequencing during the COVID-19 pandemic and to study within-host dynamics of SARS-CoV-2 (*21*).

To determine reproducibility of our assay, a subset of 12 samples determined to have high (10^4^), medium (10^3^), and low (10^2^) SARS-CoV-2 genome copy (equivalent/mL) numbers by NSP5 qRT-PCR were selected as technical replicates. ARTIC primers and Illumina adapters were trimmed, low quality bases and duplicate reads were filtered out, and mapping and variant calling were completed as described in the iVar and PrimalSeq pipeline described by Grubaugh *et al*.(*47*). Intrahost single nucleotide variants (iSNVs) were included in further analysis if they passed the Fisher’s exact test for variation above the mean error rate at that locus and had a depth of coverage at or above 100X. iSNVs were called with minor allele frequency (MAF) thresholds at 3% and 5% and compared against technical replicates (Supplemental SNS2). iSNVs detected at 3% MAF were plotted against the SARS-CoV-2 genome copy number for each sample (Supplemental SNS3A) and the number of reads mapped for each sample (Supplemental SNS3B).

To compare variation arising in the experimentally challenged mink to variation previously detected in SARS-CoV-2 circulating at mink farms, all available mink-associated SARS-CoV-2 genomes were downloaded from GISAID from 01-Jan-2020 through 22-Nov-2021. The resulting alignment of 1002 SARS-CoV-2 genome sequences included 999 with *Neovison vison* as the host species and 3 SARS-CoV-2 sequences from the genus *Mustela* (GISAID Acknowledgements, Supplemental Table SNS4).

### Serology

Sera were heat-inactivated (30 min, 56°C). After an initial 1:10 dilution of the sera, two-fold serial dilutions were prepared in DMEM. 100 TCID_50_ of SARS-CoV02 variant B.1.1.7 was added to the diluted sera. After a 1-hour incubation at 37°C and 5% CO_2_, the virus-serum mixture was added to VeroE6 cells. The cells were incubated for 6 days at 37°C and 5% CO_2_ at which time they were evaluated for CPE. The virus neutralization titer was expressed at the reciprocal value of the highest dilution of the serum that still inhibited virus replication.

### Statistical analysis

Statistical analysis was performed using GraphPad Version 8.4.3. Significance tests were performed as indicated where appropriate with reported p-values.

## Acknowledgements

We would like to thank Emmie de Wit, Brandi Williamson, Neeltje van Doremalen, Sujatha Rashid, Ranjan Mukul, Kimberly Stempe, Kent Barbian, and Sarah Anzick for their assistance with the SARS-CoV-2 isolate. The following reagent was obtained through BEI Resources, NIAID NIH: Severe Acute Respiratory Syndrome-Related Coronavirus 2, Isolate hCoV-19/England/204820464/20200, NR-54000, contributed by Bassam Hallis. We are grateful to Dr. High Hildebrant for providing our Aleution Lateral Flow test. We thank Tina Thomas, Rebecca Rosenke, and Dan Long for assistance with histology; Brian Smith, Kathy Cordova, Marissa Woods, Carl Shaia, and Taylor Saturday for technical and data assistance; Anita Mora for assistance with figures, and Rocky Mountain Veterinary Branch animal care staff for husbandry. We thank Kimmo Virtaneva and Dan Bruno for their assistance with extractions, the ARTIC protocol, and sequencing. Stacy Ricklefs assisted with the original isolate sequencing as well as the mink ACE2 sequencing. We also thank Randy Elkins, Steve Denny, and Alphie Cisar for additional support.

## Author contributions

Conceptualization: DRA, VJM

Methodology: DRA, JL, DY, JRS, CT, CC, CE, JE, SNS, VJM

Investigation: DRA, JL, JES, VAA, EH, MH, KCY, JRP, SNS, PWH, GS, DS, JVM

Visualization: DRA, JL, VAA, PWH, GS, JRS, CT, CC, NMW, SNS

Formal analysis: DRA, JL, DY, VAA, SNS, CM

Funding acquisition: VJM

Resources: JE

Supervision: VJM

Writing – Original Draft: DRA

Writing – Review and editing: DRA, SNS, JRS, VJM

## Supplementary Materials

**Supplemental Figure 1.**
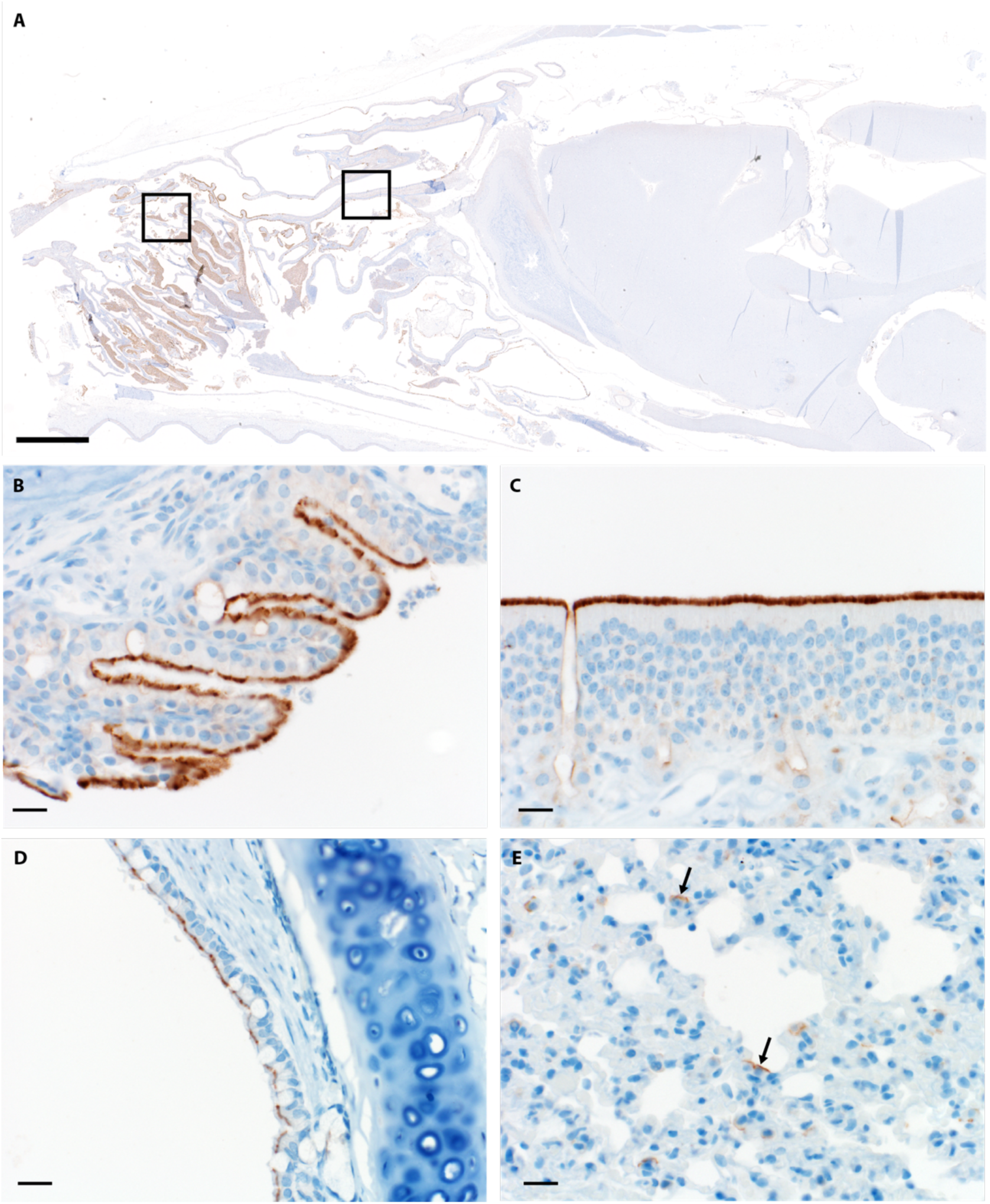
ACE2 Immunoreactivity. (A) Sagittal section of skull ACE2 immunoreactivity (brown) bar = 3mm (B) Respiratory epithelium (C) Olfactory epithelium (D) Bronchiolar epithelium (E) Type 1 pneumocytes (arrows) Bars=20um

**Supplemental Figure 2.**
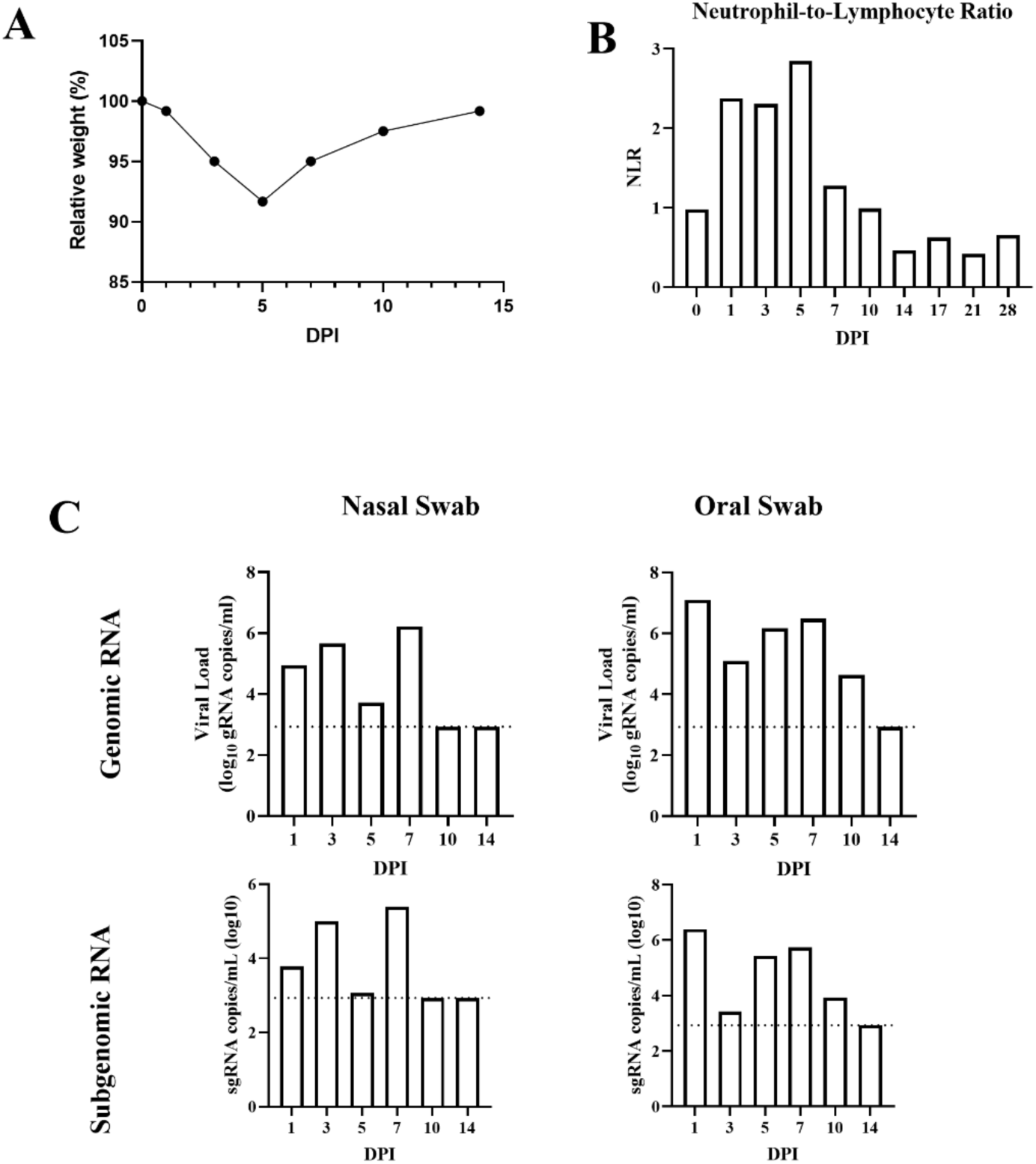
Resolution of clinical disease in the surviving animal. The surviving animal was monitored for change in relative weight (A). Weight loss was most severe on 5 DPI, after which the animal began recovery. Neutrophil-to-lymphocyte ratio was monitored over time (B), with the most severe change appreciated on 5 DPI. Nasal and oral swabs were evaluated for resolution of viral shedding through genomic and subgenomic RT-PCR (C). All swabs on 14 DPI were below the limit of detectable virus.

**Supplemental Figure 3.**
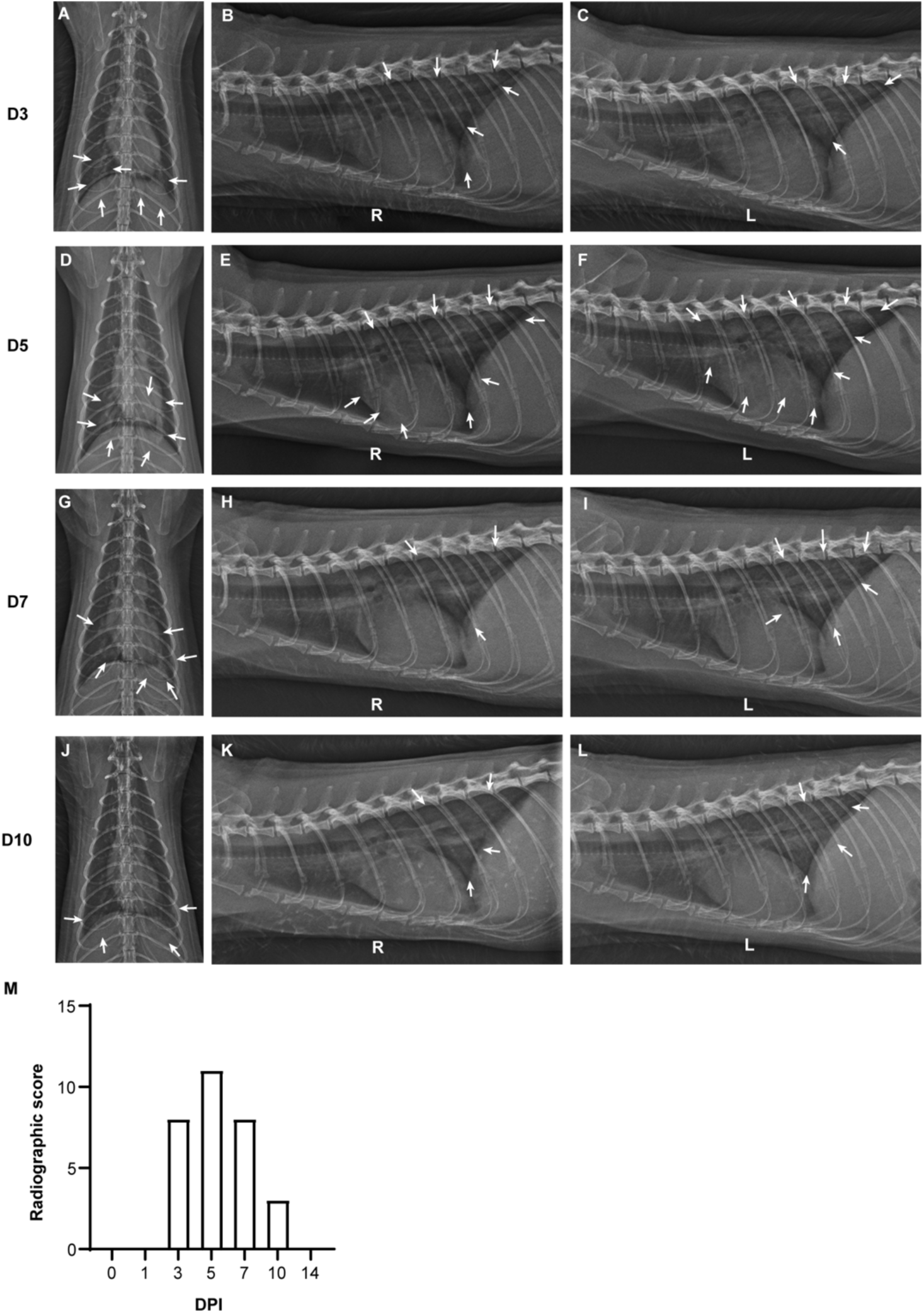
Resolution of radiological disease in the surviving animal. (A-L) Dorsoventral, right lateral, and left lateral radiographs from surviving animal on 3, 5, 7, and 10 DPI. Arrows indicate pulmonary infiltrates, first visible in the left and right caudal lung lobes at 3 DPI (A-C) with additional involvement in the caudal subsegment of the left cranial lung lobe on 5 DPI (D-F). There is mild improvement in the alveolar pattern in the left and right caudal lung lobes on 7 DPI, with resolution in the caudal subsegment of the left cranial lung lobe (G-I). The pulmonary changes continued to improve by 10 DPI, with grade 2 pulmonary disease in the right caudal lung lobe with grade 1 pulmonary disease in the left caudal lung lobe consistent with improving viral pneumonia and pneumonitis (J-L).(M) Radiographic scores for surviving animal.

**Supplemental Figure 4.**
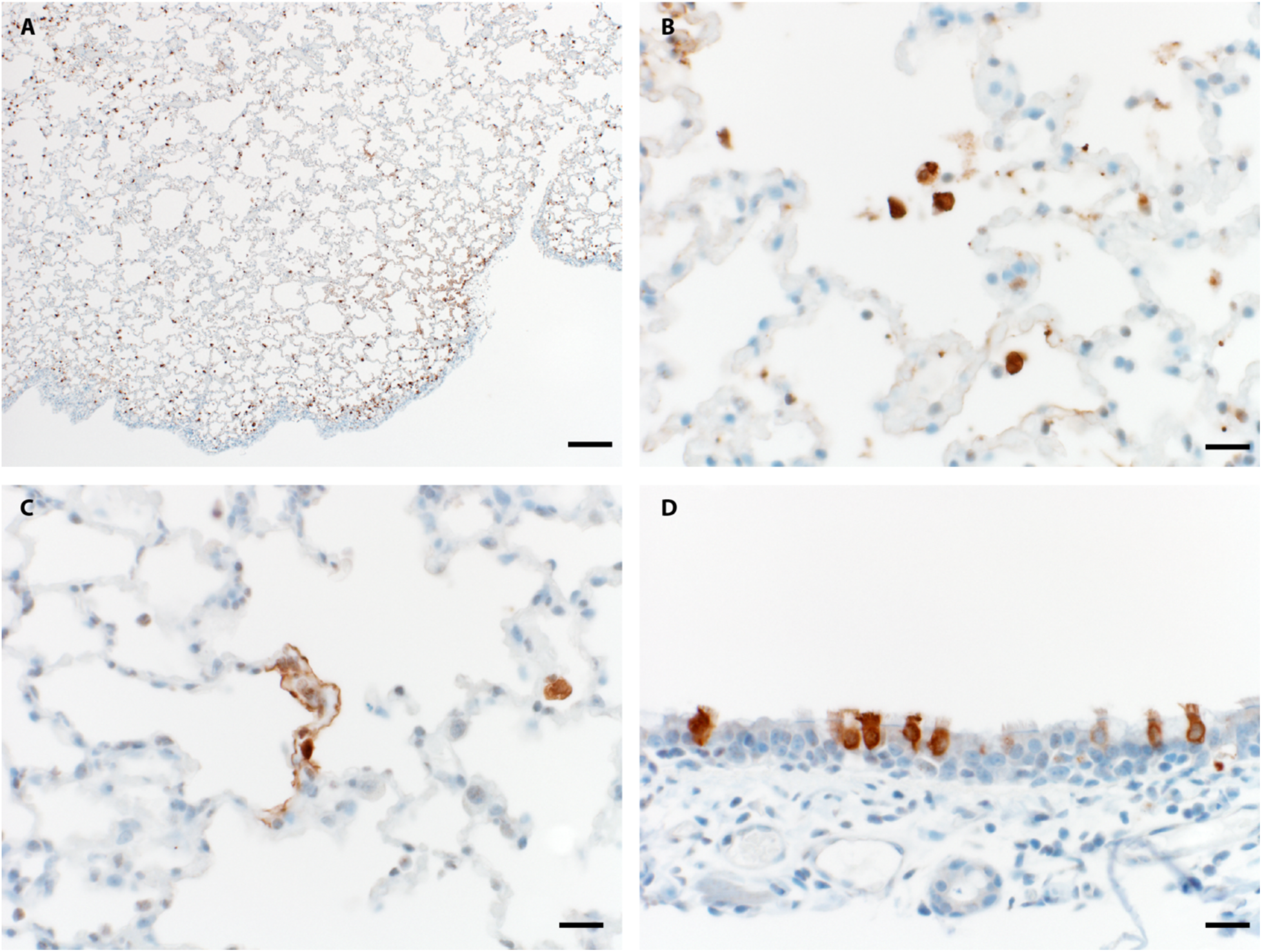
SARS-CoV-2 Pulmonary immunohistochemistry. (A) Lung: Bar=200um (B) Alveolar macrophage immunoreactivity (C) Type I & II pneumocyte immunoreactivity (D) Bronchiolar epithelium immunoreactivity (brown=immunoreactive cells) C-E Bar=20um

**Supplemental Figure 5.**
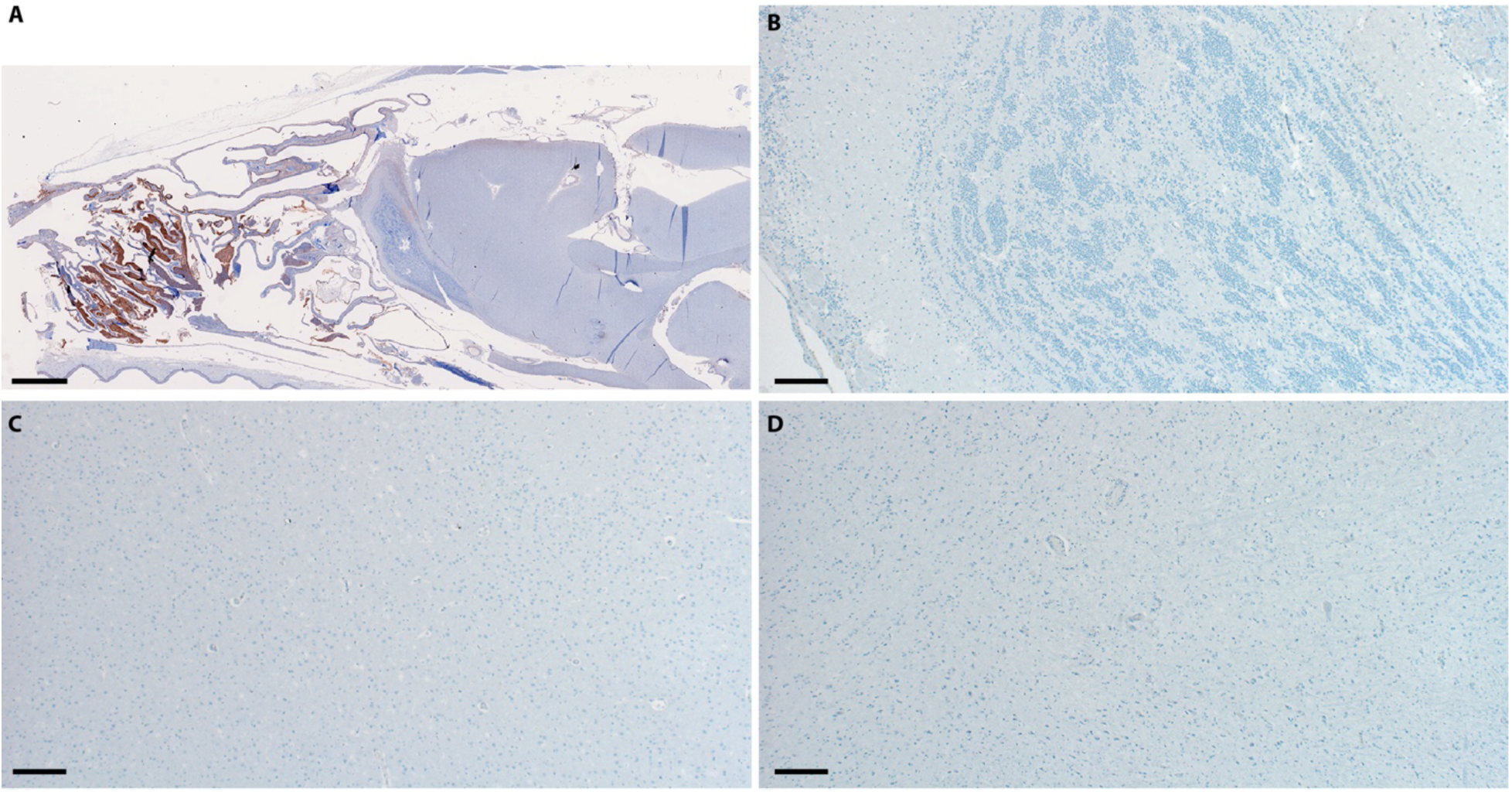
Nasal turbinate and brain SARS-CoV-2 immunohistochemistry. (A) Sagittal section of skull: Abundant nasal turbinate epithelium and exudate immunoreactivity (brown); bar = 3mm (B) Olfactory bulb: no immunoreactivity (C) Cerebral cortex: no immunoreactivity (D) Brainstem: no immunoreactivity. B-D bar = 200µm

**Supplemental Figure 6.**
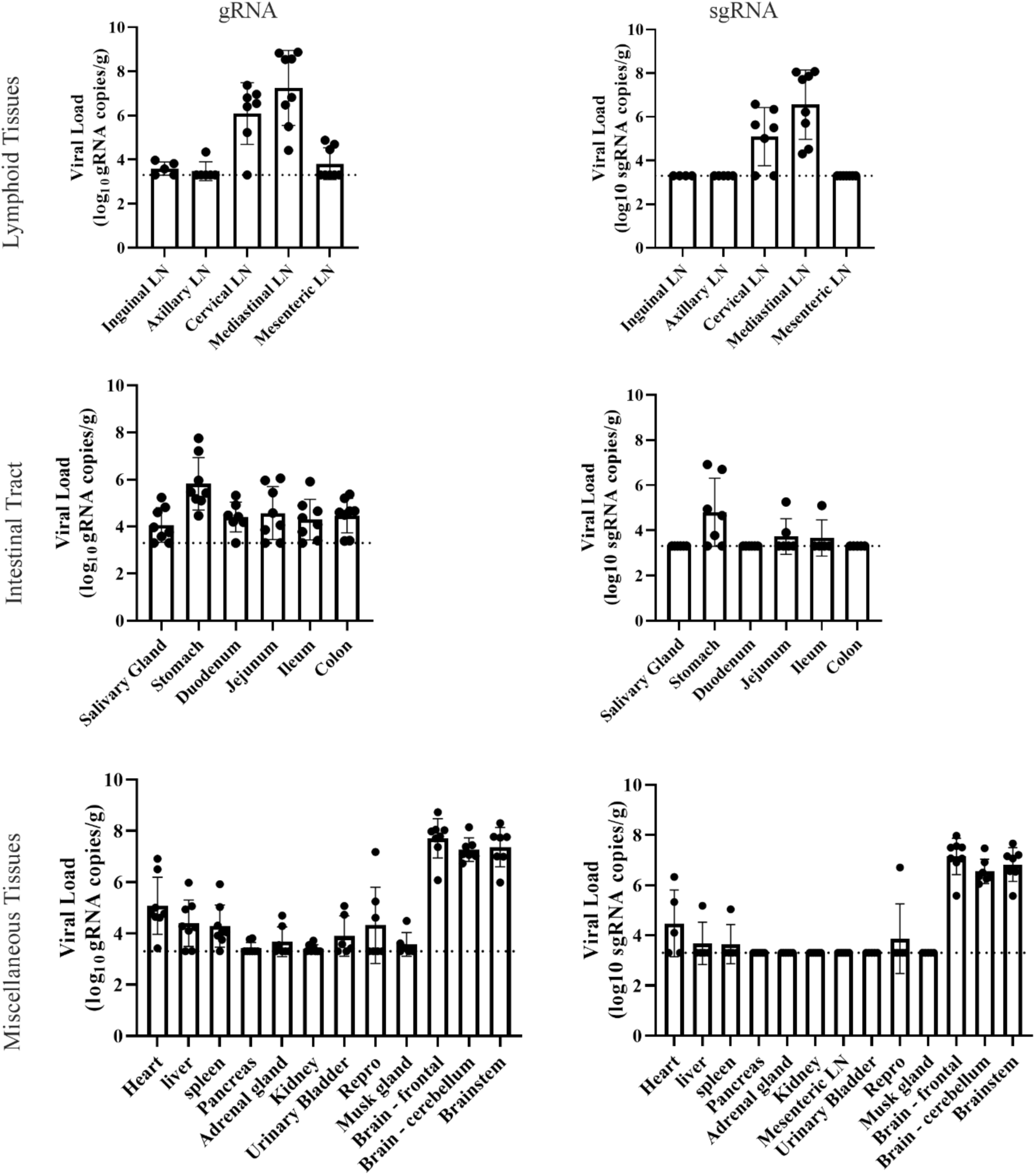
Viral burden in non-respiratory tissues. Lymphoid (A), intestinal (B), and miscellaneous (C) tissues from animals euthanized on 3 DPI were evaluated for genomic and sub-genomic RNA. Graphs depict the mean and standard deviation.

**Supplemental Figure 7.**
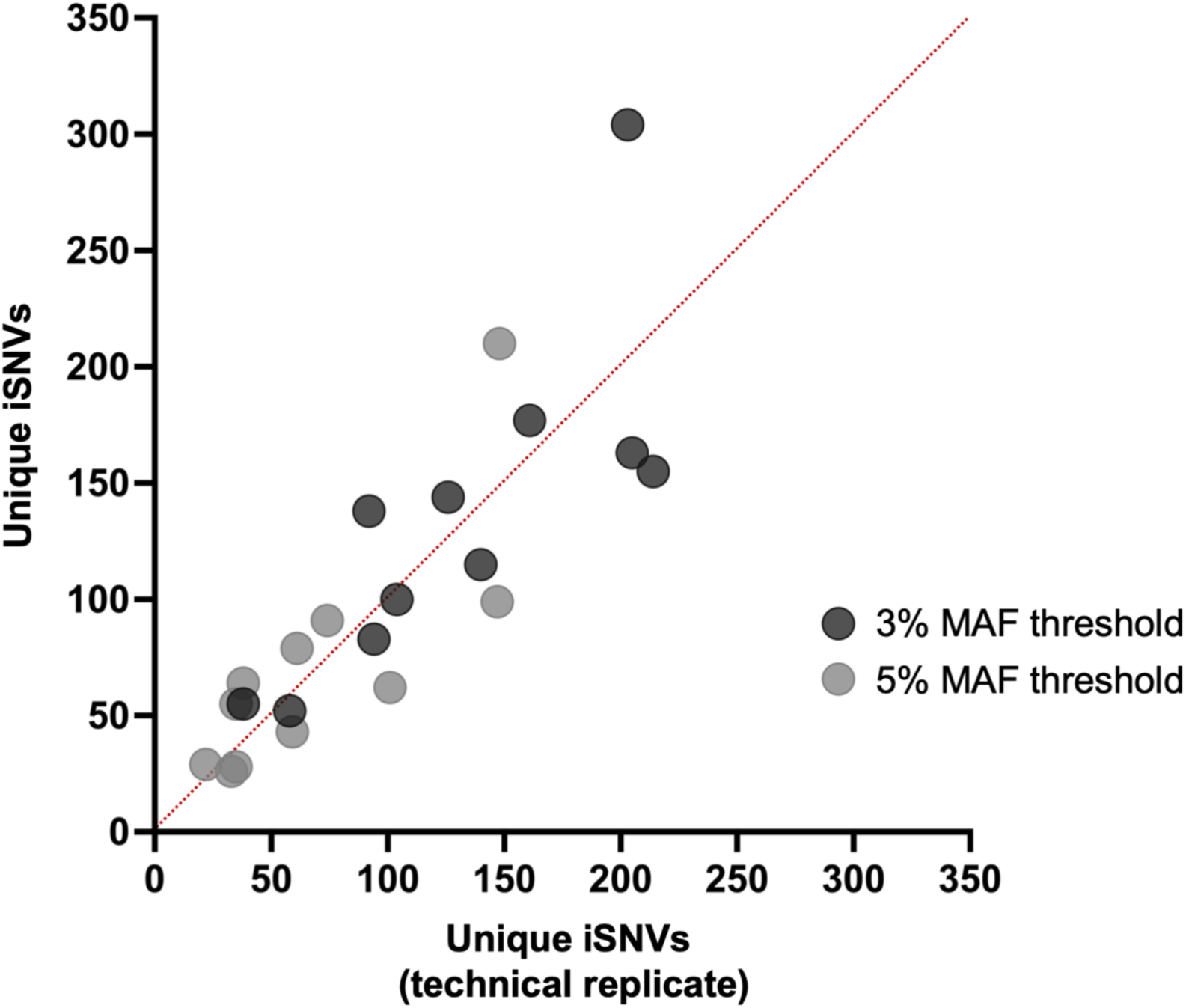
Comparison of intrahost single nucleotide variants (iSNVs) detected between technical replicates at minor allele frequencies (MAFs) of 3% and 5%. Dots along the red line indicate perfect concordance between replicates, whereas dots farther from the red line indicate less concordance between replicates.

**Supplemental Figure 8.**
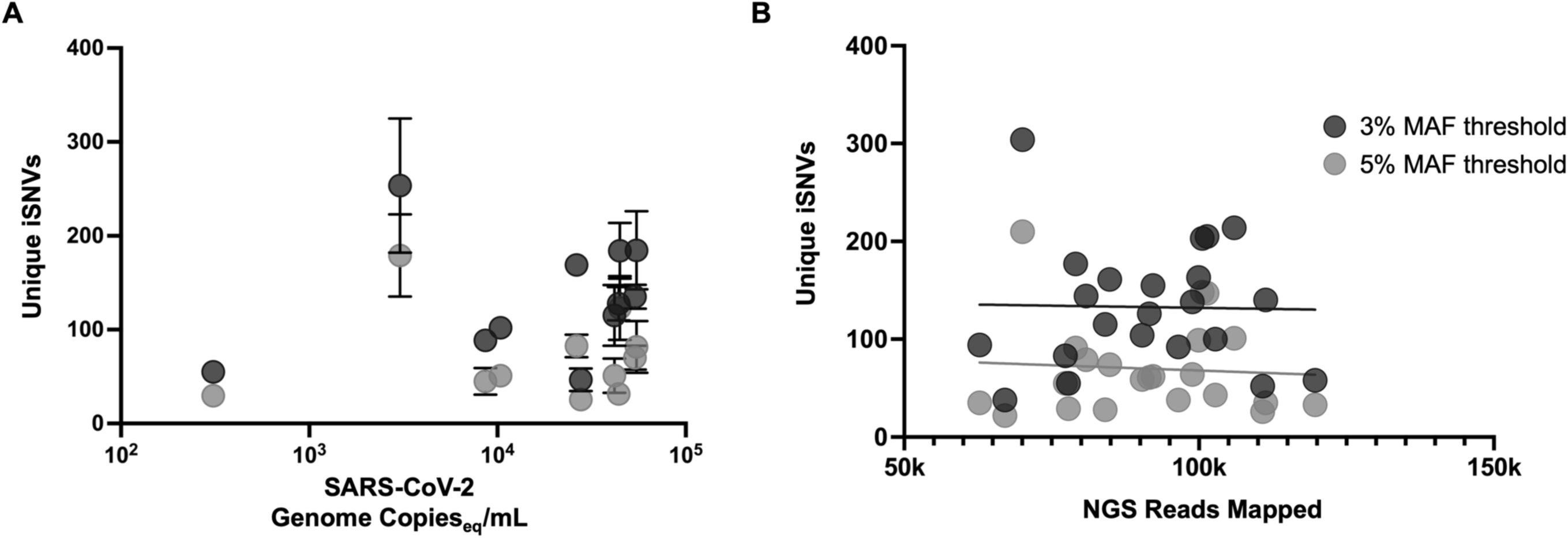
(A) Unique intrahost single nucleotide variants (iSNVs) relative to SARS-CoV-2 genome copy number or (B) number of sequenced reads mapped for each sample. Data shown for all samples with minor allele frequency thresholds of 3% and 5%.

**Supplemental Table 1.**
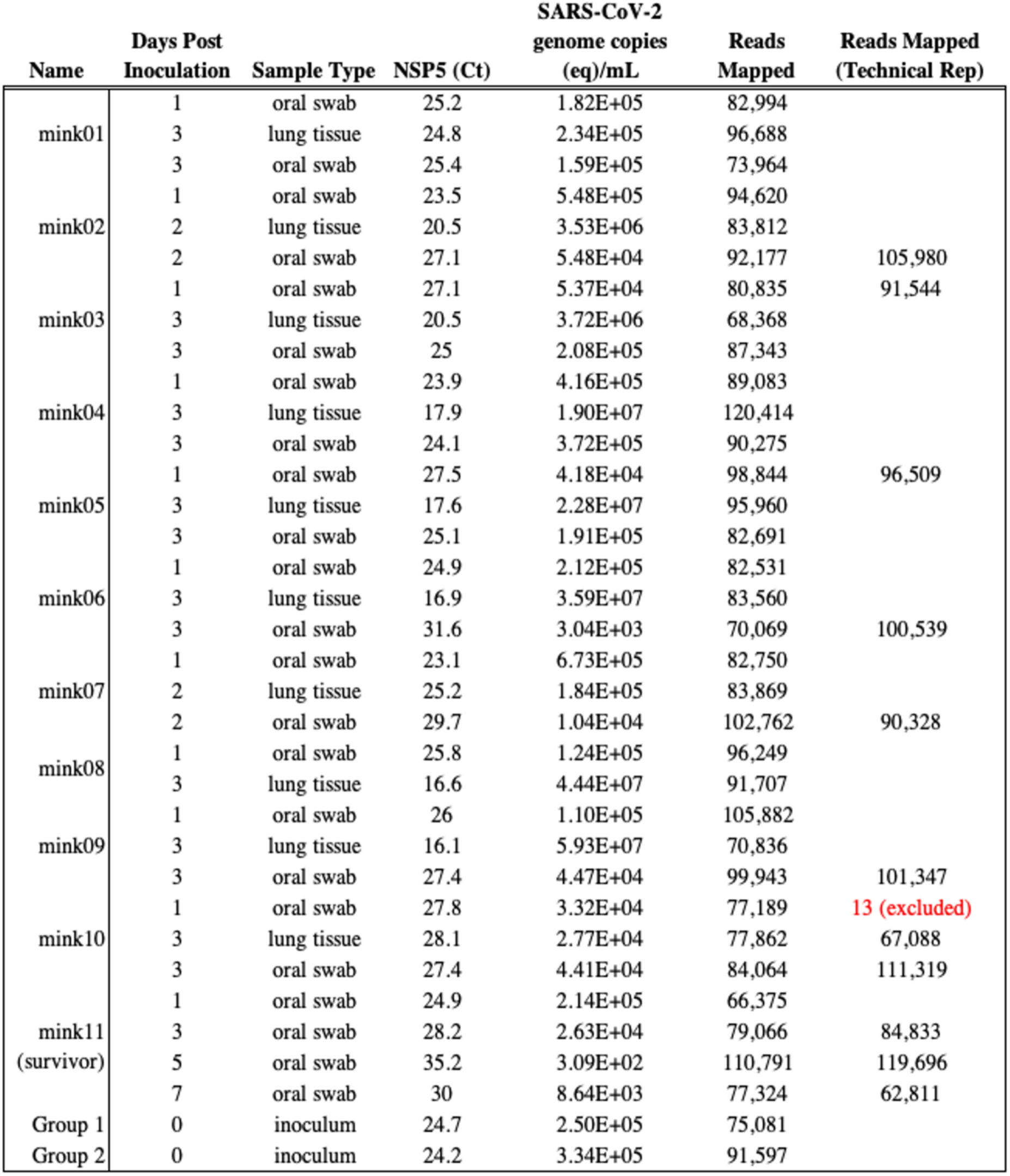
Samples from mink experimentally challenged with SARS-CoV-2 that were deep sequenced for within-host evolutionary analyses.

**Supplemental Table 2.**
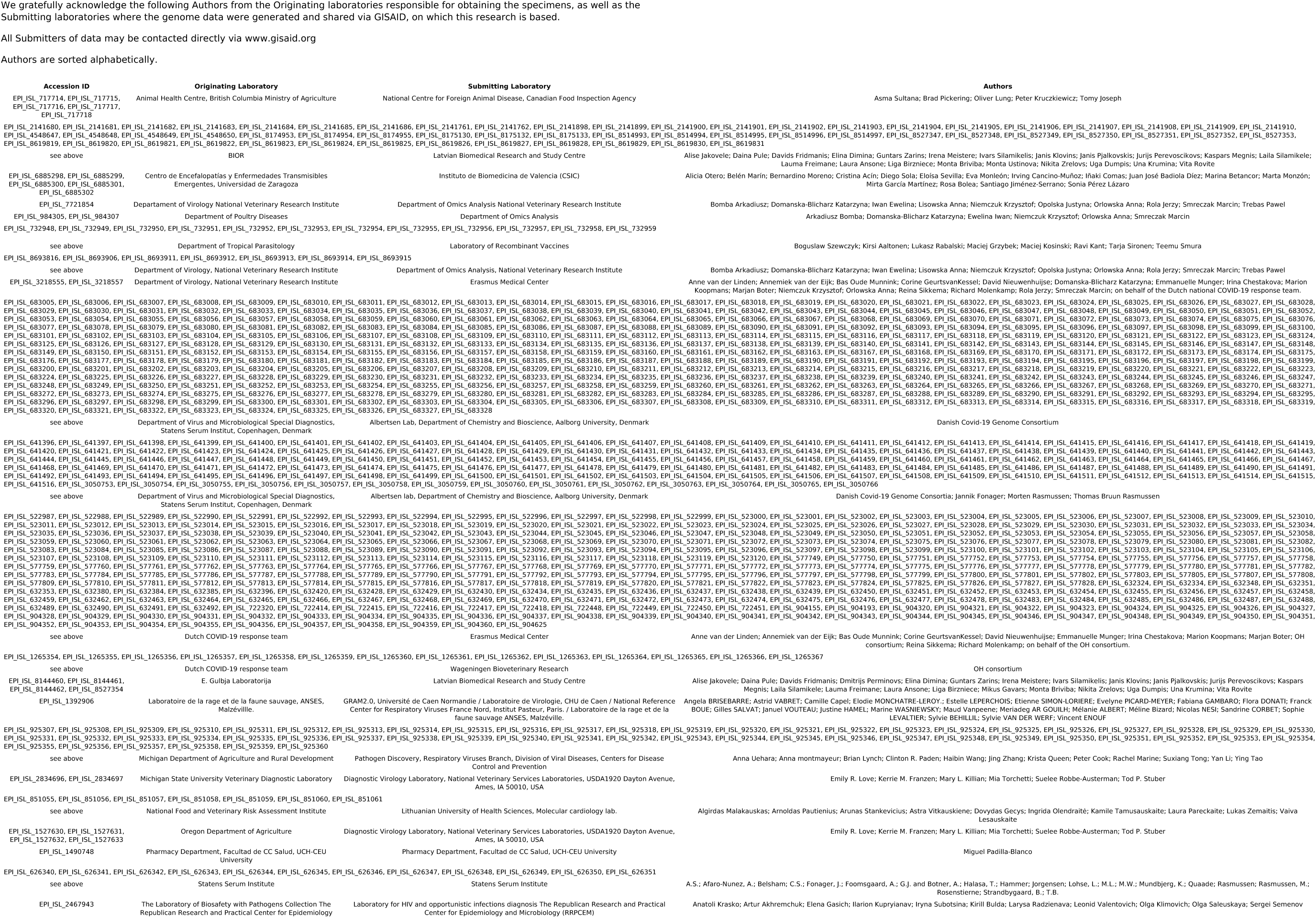
Acknowledgements for mink-associated sequences downloaded from GISAID.

**Supplementary Table 3:**
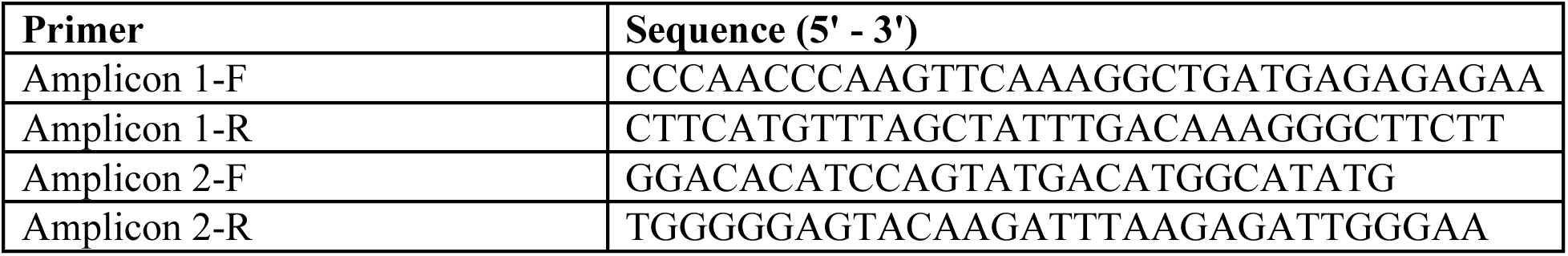
Long range PCR primers for amplification of ACE2.

